# Drought-tolerant phenotypes associated with patterns of deterministic microbiome assembly across peanut genotypes

**DOI:** 10.64898/2026.07.20.739698

**Authors:** Zoe J. Little, Deepak Shantharaj, Charles Chen, Neha Potnis

**Affiliations:** Department of Entomology and Plant Pathology, Auburn University, Auburn, Alabama AL 36849; Department of Crop, Soil and Environmental Sciences, Auburn University, Auburn, Alabama AL 36849

**Author notes:** Corresponding author: Neha Potnis. Zoe J Little – Deepak Shantharaj - Charles Chen.

**Keywords:** deterministic assembly, drought-response phenotype, endosphere, fungal communities, host filtering, host functional traits, microbial community assembly, peanut, rhizosphere

## Abstract

Plant-associated microbiomes contribute to plant health and resilience, yet the extent to which host traits shape microbiome assembly remains poorly understood despite increasing interest in leveraging them for crop performance. Here, we investigated whether drought-response phenotypes are associated with reproducible patterns of microbiome assembly across peanut genotypes under field conditions. The cultivars represented three drought-response categories: water-savers with tighter stomatal regulation, water-spenders with deeper root systems, and drought-sensitive genotypes. Bacterial and fungal communities were characterized from bulk soil, rhizosphere, and root endosphere compartments of six non-stressed peanut cultivars. Both host genotype and drought-response phenotype were associated with microbiome composition, with phenotype-associated patterns remaining detectable across multiple genetic backgrounds. Unexpectedly, the strongest phenotype-associated differences occurred in bulk soil communities, suggesting plant-mediated effects extending beyond the immediate root zone. Community differences were driven primarily by shifts in the relative abundance of existing taxa rather than turnover of distinct microbial lineages. Fungal communities responded more strongly to host phenotype than bacterial communities, with water-spender genotypes supporting greater fungal diversity and uniquely enriched taxa in the rhizosphere and endosphere. Neutral community modeling indicated stronger deterministic filtering of fungi than bacteria. Together, these findings demonstrate that drought-response phenotypes shape reproducible microbiome variation before stress exposure.

**Highlight:** This study investigates the potential for host phenotype-associated drivers of microbiome assembly in drought-tolerant peanut cultivars that represented different physiological mechanisms for drought tolerance.

## Introduction

Peanut is one of the most widely cultivated legumes globally (Wagner, 2021). They are a crucial part of agricultural ecosystems in tropical and subtropical regions of Asia and Africa, where they largely contribute to food security and soil fertility (due to biological mechanisms such as nitrogen-fixation) (Wagner, 2021). A large percentage of peanut production occurs in semi-arid or seasonal rainfall zone, where drought episodes in regions reliant on these rain-fed agricultural systems cause devastating yield loss (Oyserman et al., 2022). Due to this, drought has been identified as a dominant threat to the peanut industry where it contributes to roughly 50 million USD in losses annually (Pokhrel et al., 2025). As the severity and duration of droughts are expected to increase alongside rapidly changing environmental conditions, new strategies are being developed to increase crop resilience to drought and other abiotic stressors alongside other mitigation efforts like enhancing water use efficiency and irrigation redesigns.

Plant hosts exhibit intimate relationships with complex networks of bacteria, archaea, fungi, and viruses that are collectively referred to as the plant microbiome. While it is well known that microorganisms help to maintain fundamental aspects of all life, the plant microbiome has gained increased attention for its role in modulating plant health (Turner et al., 2013). Plant microbes are known to assist in nutrient acquisition, protection against biotic and abiotic stressors, and overall growth promotion of the host (Yusuf et al. 2025). Different areas of a plant, or “plant compartments”, provide specialized environments for microorganisms that are uniquely enriched depending on the requirements of the host (Rossmann et al., 2017). Bulk soil plant compartment refers to the area of soil around a plant and this area acts as a pool of potential microbiota for other areas of the plant (Ling et al., 2022). Microbial community members from this pool are selectively recruited for colonization into the root structure (often referred to as the rhizosphere) and are known to interact synergistically with the host to contribute to plant growth, mediation of phytohormone production, and priming of plant immune defenses (Kumar et al., 2018). Beyond the rhizosphere, the root endosphere refers to the space within the root tissue of a plant that is colonized by a highly specific group of microbiota, either by horizontal or vertical transmission (Lengrand et al., 2024). In terms of plant-mediated microbiome assembly, selection of and colonization by specific microbes moves in an increasingly selective spatial and biochemical gradient from bulk soil to root endosphere (Kalumbilo et al., 2026). A combination of factors such as plant species, genotypes, soil types, abiotic stress, season, cultural practices, and predation can determine how plant microbial communities are assembled among specialized compartments (Guo et al., 2023).

There is increasing evidence that host genotype shapes the composition and assembly of the plant-associated microbiome, with downstream consequences for host performance under both biotic and abiotic stress. Model systems like *Arabidopsis* have shown that bacteria belonging to phylum *Actinobacteria* differentially colonize roots in response to cues derived from metabolically active host cells (Bulgarelli et al., 2012). In crop systems, similar genotype-dependent effects have been observed. In wheat, disease resistant cultivars were consistently associated with beneficial early-colonizing rhizosphere taxa, contributing to a distinct metabolite profile that supported viral disease suppression (C. Wu et al., 2025). In maize, host genotype explained a significant amount of variation in rhizosphere microbiota even after controlling for environmental and technical factors (Peiffer et al., 2013). These genotype effects are an indirect byproduct of genetically determined phenotypic traits that modify root exudate-mediated selection processes, resulting in the regulation of microbial recruitment across rhizosphere and endosphere compartments (Bouwmeester et al., 2025).

As previously mentioned, a central mechanistic driver of microbiome assembly is genotype-to-phenotype expression of root exudate profiles, including primary and secondary metabolites such as carbohydrates, amino acids, phenolics, flavonoids, and auxins (L. Wu et al., 2023; Yang et al., 2024). These compounds establish chemical gradients that drive microbial movement toward or away from host root tissues, and colonization at the root-soil interface (Jing et al., 2023). While a host’s genotype determines biosynthetic capacity for these metabolites, it is ultimately the host phenotype, shaped by developmental state and environmental context, that defines the quantity and composition of exudates, and consequently the microbial communities that assemble (Taiz et al., 2015). Beyond exudates, phenotypic traits such as root architecture, and immune signaling further act as filters that drive microbial assembly of symbiotic microbes, not only in rhizosphere communities but also the subset of microbes that successfully transition into the endosphere (Araujo et al., 2025). This hierarchical selection results in the enrichment of specific microbial communities with the host that can functionally integrate with host physiology and support overall host health (Brachi et al., 2022; Wagner, 2021). Thus, these microbiomes are recognized not merely as passive assemblages but as active contributors to host fitness, providing microbiome-mediated protection through mechanisms such as pathogen suppression, immune priming, and enhanced tolerance to abiotic stresses (González & Elena, 2021; Singh et al., 2023). Evidence for this functional linkage is emerging across host systems. For example, in oat (*Avena sativa*) and wheat (*Triticum aestivum*), host genotype influences microbiome assembly under water stress, likely through differential stress-induced shifts in metabolite profiles among different host genotypes that selectively enrich drought-adaptive microbes (Guo et al., 2023).

These recruited microbiomes can, in turn, enhance host resilience by modulating stress-responsive pathways, improving nutrient acquisition, and stabilizing root function under stress (Fadiji et al., 2023; Hawkes et al., 2020; Pérez-Izquierdo et al., 2019). Therefore, we deduce that abiotic or biotic stress tolerance in plants can be viewed as emergent property of plant-microbiome holobiont.

Auburn University’s peanut breeding program has shifted from selection solely for high yield toward integrating physiological traits associated with drought tolerance (Zhang et al., 2022). This transition has been guided by analogous efforts in other crops, for example, improved water use efficiency (WUE) has enhanced drought resilience in wheat, while increased biological nitrogen fixation (BNF) has supported yield stability in soybean under water-limited conditions (Condon et al., 2002, 2004; Sinclair et al., 2007; Zhang et al., 2022). Besides agronomical traits, evaluation of peanut drought tolerance at Auburn University has extended to the traits that include denser root systems that set deeper into the soil, transpiration efficiency (TE) while (sustaining photosynthetic capabilities), improved biological nitrogen fixation (BNF), harvest index, and overall biomass accumulation (Zhang, 2021). Deeper root systems and improved TE are identified as the most studied traits of drought tolerance in peanut. This is due to their relevance in addressing water availability which remain one of the largest abiotic stressors influencing peanut yield (Zhang, 2021).

Variation in TE among peanut genotypes is largely mediated through stomatal regulation (Zhang et al., 2022). Under drought conditions, tighter stomatal control reduces transpiration but also limits carbon-di-oxide diffusion to photosynthetic sites (Tardieu et al., 2018). Genotypes that adopt this conservative water-use strategy are classified as “isohydric” or “water-saver” genotypes (Zhang et al., 2022). Physiological mechanisms of water-saver genotypes are associated with chemical and hydraulic signaling which are thought to be influenced by gradients of the phytohormone abscisic acid (ABA) (Tardieu et al., 2018). Notably, root-associated microbiomes are known to play a crucial role in driving change in root exudate and phytohormone production suggesting a potential microbial contribution to water-saver phenotypes (Sharma et al., 2025; Tardieu et al., 2018).

However, the effectiveness of water-saver strategies is context dependent, with the most benefits seen in dry seasonal environments with longer periods of intense drought. In regions such as the Southeastern United States, where drought is typically intermittent and rainfall remains possible, selecting genotypes with highly regulated stomatal control may lead to decreased yield output (Zhang et al., 2022). In these environments, these “water-spender” genotypes that utilize deeper root systems with dense root architecture may be more advantageous to combat issues of water availability by i) reaching larger areas of sporadically moist soil and ii) harboring the required biomass to retain moisture (Zhang et al., 2022). Given the critical role of the rhizosphere microbiome in regulating root function, including phytohormone modulation, nutrient acquisition, and stress priming, understanding microbial community composition in water-spender genotypes is essential for evaluating their functional potential under variable drought conditions. Knowing the crucial role that rhizosphere microbiota has on the functionality and efficiency of root systems, investigation into the microbial community that is present in water-spender peanut genotypes are particularly necessary. Water-saver and water-spender phenotypes represent contrasting drought-adaptive strategies that provide a useful framework for investigating the biological mechanisms underlying drought resilience.

We therefore hypothesized that peanut genotypes exhibiting distinct drought response strategies would harbor unique root-associated microbial communities. Because microbiome composition can be influenced by host genetic background, we included two genotypes within each drought-response category (water-saver, water-spender, and drought sensitive) to minimize the influence of any single genotype on observed community patterns. The water-saver genotypes (Line 8 and AU-16-28) share a common parental lineage but originated from different breeding crosses, whereas the water-spender genotypes (AU-NPL-17 and PI-502120) represent more genetically distinct backgrounds with PI-502120 being a drought tolerant Peruvian landrace accession. We reasoned that microbial taxa or community features consistently enriched across genotypes exhibiting the same drought-response phenotype would provide preliminary evidence for phenotype-associated microbial assembly, whereas taxa restricted to individual genotypes would more likely reflect genotype-specific effects. This framework enabled us to evaluate whether microbiome assembly patterns are linked to drought-response categories rather than individual host genotypes. As resistant plant breeding moves toward integrated forms of plant resistance (genetically resistant genotypes paired with sustainable, non-chemical pest and pathogen controls), it is becoming mandatory to acknowledge the plant-associated microbiome’s role in determining the success of resistance traits.

## Methods

In this study, six genotypes of Line 8, AU-NPL 17, AU16-28, PI502120, PI390428, and AP-3 were planted in Auburn University Wiregrass Research and Education Station (AAES) at Headland, AL. Six peanut genotypes were selected from three unique phenotypic classifications of drought tolerance (water-saver, water-spender, and sensitive) from Auburn University’s peanut breeding program (Table 1). The experiment plots were two-row plot of 6.10 meters long and 91 cm space between row and the experiment design was randomized complete block design with 3 replications. The seeds were planted on May 22, 2024. Plants were sampled in early October 2024, 135 days after planting, which marked the final harvest period of the growing season. From each plot, three plants were randomly selected for sampling from the first genotype; these randomly selected locations determined sampling for the other five genotypes (Fig. S3). The samples present in this study are from 6 different genotypes spanning across 3 different host genotype categories watered under rain-fed and irrigated systems. Each sample was further separated by plant compartment, bulk soil, rhizosphere soil, and root endosphere tissue for microbiome profiling. In total, there were 56 samples sequenced for this study including both bacterial and fungal amplicon sequencing. There are positive controls in the form of Zymo community standards and Zymo DNA samples (*ZymoBIOMICS Microbial Community DNA Standard*, n.d.) and negative controls in the form of buffer extractions.

**Table 1:**
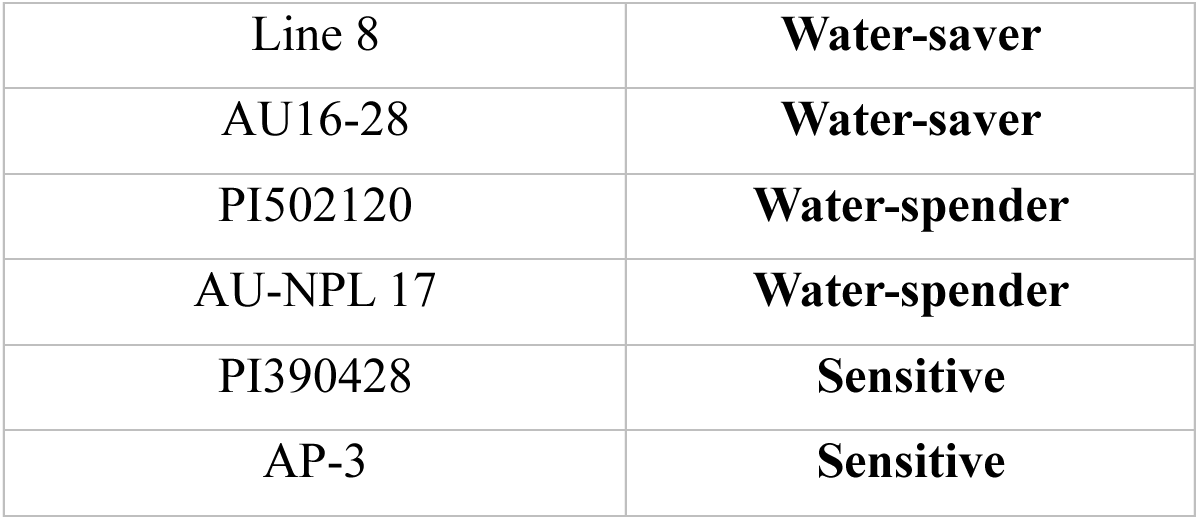
genotype-to-water habit classifications of samples included in study.

|  |  |
| --- | --- |
| Line 8 | **Water-saver** |
| AU16-28 | **Water-saver** |
| PI502120 | **Water-spender** |
| AU-NPL 17 | **Water-spender** |
| PI390428 | **Sensitive** |
| AP-3 | **Sensitive** |

### Sample Preparation and DNA Extraction

To collect samples for each compartment, peanut plants were carefully loosened and uprooted from the soil and the intact root system was cut at the soil line and stored in sealable plastic bags at 4℃ for transport. To collect bulk soil, samples were taken from the remaining soil that separated from the plant while lifting it from the ground. This soil was roughly 4 inches deep and was collected in 5ml Eppendorf tubes to be stored at 4℃ until DNA extraction. For the separation of rhizosphere soil from the intact root tissue, a sample of root cutting was placed in a 50mL falcon tube and 30-45mL of 10mM NaCl solution was added to completely cover the root tissue. Root cuttings were sonicated using the VEVOR Digital Ultrasonic Cleaner (Model: MH-040S) for 5 minutes and vortexed. If any rhizosphere soil remained attached to the root tissue, another NaCl wash was performed, and the resulting pellets were combined prior to DNA extraction. After cleaning the root tissue fully, it was removed and placed in a Ziploc bag between Kimwipes to dry overnight. The remaining NaCl solution was centrifuged at 12,000 rcf for 10 min and, after decanting the supernatant, the resulting rhizosphere pellet was stored at -20 ℃ until DNA extraction. To perform the extractions, total microbial DNA was extracted from 0.25g of bulk soil samples using the Qiagen DNeasy PowerSoil Pro Kit with modifications. A 1mL aliquot of the rhizosphere pellet was transferred to a 2 mL Eppendorf tube and DNA was extracted using modified a modified CTAB protocol (see a detailed extraction protocol on GitHub https://github.com/Potnislab/peanut_genotype_microbiome_manuscript). To prepare the clean root tissue for endophytic DNA extraction, a 0.5 – 1g sample of the frozen root tissue was finely cut up with sterile scissors and placed in a 2mL Eppendorf tube. Sterile BB beads were added to the tubes and were homogenized using the BioSpec Mini Bead Beater 96 (Product No. 1001). After homogenizing the root tissue, the BB beads were removed with a magnet, and the total microbial DNA was extracted using the same modified CTAB protocol as the rhizosphere soil samples. The DNA samples were quantified using the Qubit 2.0 Fluorometer (ThermoFisher Scientific, 2011).

To sequence the bacterial and fungal taxa present in the samples, quantified DNA samples were sent to University of Connecticut’s Microbial Analysis, Resources and Services (MARS) facility for high-throughput Illumina MiSeq 2x250 sequencing. Conditions during PCR were as follows: initial denaturation at 95°C for 2 min, 30 cycles at 95°C for 20 s, 75°C for 10 s (the addition of PNA clamps to the root endosphere samples, clamping temperature), 55°C for 20 s, 72°C for 1 min, and a final extension at 72°C for 10 min. Two PCR blanks were included. Next, the amplicons were purified using Ampure XP (Coulter Beckman, n.d.) and the DNA concentration of the purified samples was quantified using a Qubit 3.0 fluorometer (ThermoFisher Scientific, 2014). These DNA concentrations were used to pool samples and blanks in equimolar concentrations, resulting in a library. The amplicon library was further purified by loading it on a 0.8% (mass/vol) agarose gel and extracting bands of approximately 380 bp with the Nucleospin Gel and PCR Clean-up kit (Macherey-Nagel, n.d.). For each round of PCR containing the root endopshere tissue, 0.25 μL mPNA and 0.25 μL pPNA were added to block potential host DNA amplification. V4 regions were amplified using 515F and 806R Illumina adapters and ITS regions were amplified with ITS2.

### Data Preparation

After receiving raw FASTQ files back from the MARS sequencing facility, raw paired-end Illumina reads were downloaded and input into Alabama ASAX HPC supercomputer for data processing. Adapter trimming was performed and primers were removed with BBduk (Bushnell, 2014) and bases with a Phred quality score of <20 were removed. Before quality filtering and assessments were performed, the bulk soil, rhizosphere, and root endosphere reads were separated for easier processing in HPC. Quality checks were performed using FastQC (Andrews, n.d.) where the quality of bases was checked and singletons were removed and validated which also assessed successful removal of primer sequences. Error models were generated and quality was assessed using DADA2 v1.40.0 (Callahan et al., 2016). Prior to ASV inference, the bulk soil, rhizosphere, and root endosphere samples were re-merged and processed together until later downstream statistical analyses of plant compartments. ASV inference was performed using DADA2; paired-end reads were imported and denoised, dereplicated, and re-replicated. The reads were then merged and chimeric sequences were removed. After ASV inference, taxonomic classification was assigned using the DECIPHER package and DADA2. For the bacterial dataset, the SILVA SSU r138 (2019) database was used as the taxonomic reference. For the fungal dataset, the UNITE sh_general_release_dynamic (19.02.2025) dataset was used. Negative (buffer) controls from all extraction protocols were sequenced to assess potential contamination and were excluded from downstream analyses after the lack of amplified microbial reads was confirmed. For the 16S rRNA dataset, sequences assigned to host mitochondria and non-target eukaryotic taxa were removed. Samples containing fewer than 1,000 sequencing reads were excluded to ensure consistent sequencing depth for downstream analyses. As a result of this minimum sequencing threshold, two endosphere samples were excluded prior to analysis in both datasets. This resulted in a final replication of 2–3 samples per genotype and 5–6 samples per phenotype, which is a minor imbalance that should be considered when interpreting genotype-level comparisons of endosphere communities. Due to the hierarchical structure of genotypes being categorized within phenotypes, we opted to apply a stratified PERMANOVA analysis to avoid reducing the statistical power of phenotype groupings further and interpreted endosphere effects as exploratory. The remaining samples were then rarefied to an even sequencing depth using the phyloseq package.

Alpha diversity (diversity within a sample) of the bacterial and fungal communities was calculated from amplicon sequence variant (ASV) tables using the estimate_richness function in phyloseq package v1.56.0 in R (McMurdie & Holmes, 2013). These diversity metrics were assessed through Shannon Index (H) and Chao 1 richness using the phyloseq package. Beta diversity (dissimilarity between samples) was assessed through the vegan package v2.7.5 (Oksanen et al., 2026). Beta diversity was assessed using both phylogenetic and non-phylogenetic distance metrics to capture the phylogenetic nuance of any potential community variation. Phylogenetic beta diversity was calculated using unweighted UniFrac distances, which takes phylogenetic information between taxa into account while incorporating presence–absence scores (Fukuyama et al., 2012). In addition, non-phylogenetic compositional dissimilarities were calculated using Bray–Curtis dissimilarity to assess abundance-based diversity dissimilarities (IMPACTT investigators, 2022). Principal coordinates analysis (PCoA) was performed with the vegan package to visualize patterns of community dissimilarity patterns seen in beta diversity analyses. To assess how specific host variables affected microbial assembly patterns, a constrained principal coordinates analysis (CAP) was performed using unweighted Unifrac and Bray-Curtis distance metrics. The ordination was computed by using the model formula: Host_Phenotype + Host_Genotype + Replicate. Prior to ordination and beta-diversity analyses, phylogenetic trees associated with each dataset were assessed to ensure consistent rooting. For each rarefied phyloseq object that was separated earlier in the analysis (bulk soil, rhizosphere, and endosphere compartment samples), the phylogenetic tree was midpoint-rooted using phyloseq package. All further statistical analyses and visualizations were conducted in R, and a fixed random seed was set prior to the phyloseq rooting step to ensure reproducibility.

### Statistical Analysis

Normality of the data was assessed using Shapiro–Wilk tests, and because at least one metric deviated from normality across datasets, Kruskal–Wallis tests followed by Dunn’s post hoc comparisons with a Benjamini–Hochberg correction was used to assess differences among host genotypes, host phenotypes, and compartments. Differences between host phenotype, host genotype, and plant compartments were evaluated using permutational multivariate analysis of variance (PERMANOVA), 9999 permutations) which was conducted using the vegan package in R (Anderson, 2001). To confirm the validity of significant PERMANOVA results, homogeneity of multivariate dispersion was assessed using the betadisper function to determine whether observed differences among groups were due to true differences in centroid location (Anderson, 2017).

The 15 most abundant genera were identified by pruning the dataset used in this portion of the analysis so that it only contained reads from the top 15 genera. Stacked bar plots were constructed in ggplot2 (Wickham, 2016) with genera ordered consistently across bars in ascending order of mean relative abundance so that variations in the most abundant taxa could be visually tracked across host phenotype groupings. For the ITS endosphere dataset, one sample (1 replicate of genotype AP3) was excluded in this analysis because it contained zero reads after the rarefaction step, and thus all inferences made from the fungal endosphere top 15 genera analysis were interpreted in context of this sampling power.

Differential abundance analyses were performed separately for bulk soil, rhizosphere, and root endosphere microbial communities using the DESeq2 v1.52.0 package in R (Love et al., 2014). Unrarefied ASV count tables derived from phyloseq objects were used as input and host phenotype was used as the explanatory variable. In this study, DESeq2 was used to as a model to calculate the overall difference in taxa enrichment between host phenotype groups using generalized linear models with Wald significance tests. P-values were adjusted for multiple testing using standard Benjamini–Hochberg false discovery rate (FDR) procedure. Due to sparsity of taxa in the root endosphere because of sequencing quality control, size factors were estimated using the “poscounts” to exclude taxa that only contained zero reads. For all compartments and phenotype groupings, taxa with adjusted p-values below 0.05 were considered significantly differentially abundant (Love et al., 2014). The Sloan neutral community model (NCM) was used to evaluate the potential importance of stochastic processes to microbial community assembly across all three plant compartments (Sloan et al., 2006).

## Results

### Host Phenotype Influences Richness and Evenness of Bacterial Communities in Bulk Soil and Fungal Communities in Rhizosphere and Root Endosphere

Alt text: Two alpha diversity box plot panels representing the community diversity changes between host phenotype groupings. There is one panel for bacterial (16S) communities, showing Shannon diversity and Chao 1 diversity changes between each host phenotype and are separated for bulk soil, rhizosphere, and root endosphere. The next panel shows identical information for fungal communities.

In the bulk soil compartment, bacterial alpha diversity differed significantly among host phenotype groups for both Shannon diversity (Kruskal–Wallis test, χ² = 15.158, df = 2, p = 0.001, Fig. 1; Table S1.1) and Chao 1 richness (KW, χ² = 15.252, df = 2, p = 0.001; Table S1.1) Dunn’s post doc comparisons revealed that these differences were primarily driven by contrasts between drought sensitive and water-spender phenotypes for both diversity metrics (p =0.0002, Table. S1.2). At the genotype level, bacterial community diversity also varied significantly, as reflected by differences in both Shannon diversity (KW, χ² = 15.924, df = 5, p = 0.007; Table S1.1) and Chao1 richness (KW, χ² = 16.229, df = 5, p = 0.006, Table S1.1). Notably, the water-spender genotypes, AU-NPL-17 and PI502120, differed significantly from the drought-sensitive genotypes, PI-390428 and AP-3, respectively (Table S1.3), suggesting that the observed diversity patterns were consistent with host drought response phenotype classifications. In contrast, fungal alpha diversity in bulk soil did not differ significantly among host phenotypes or genotypes (Table S4.1).

**Figure 1:**
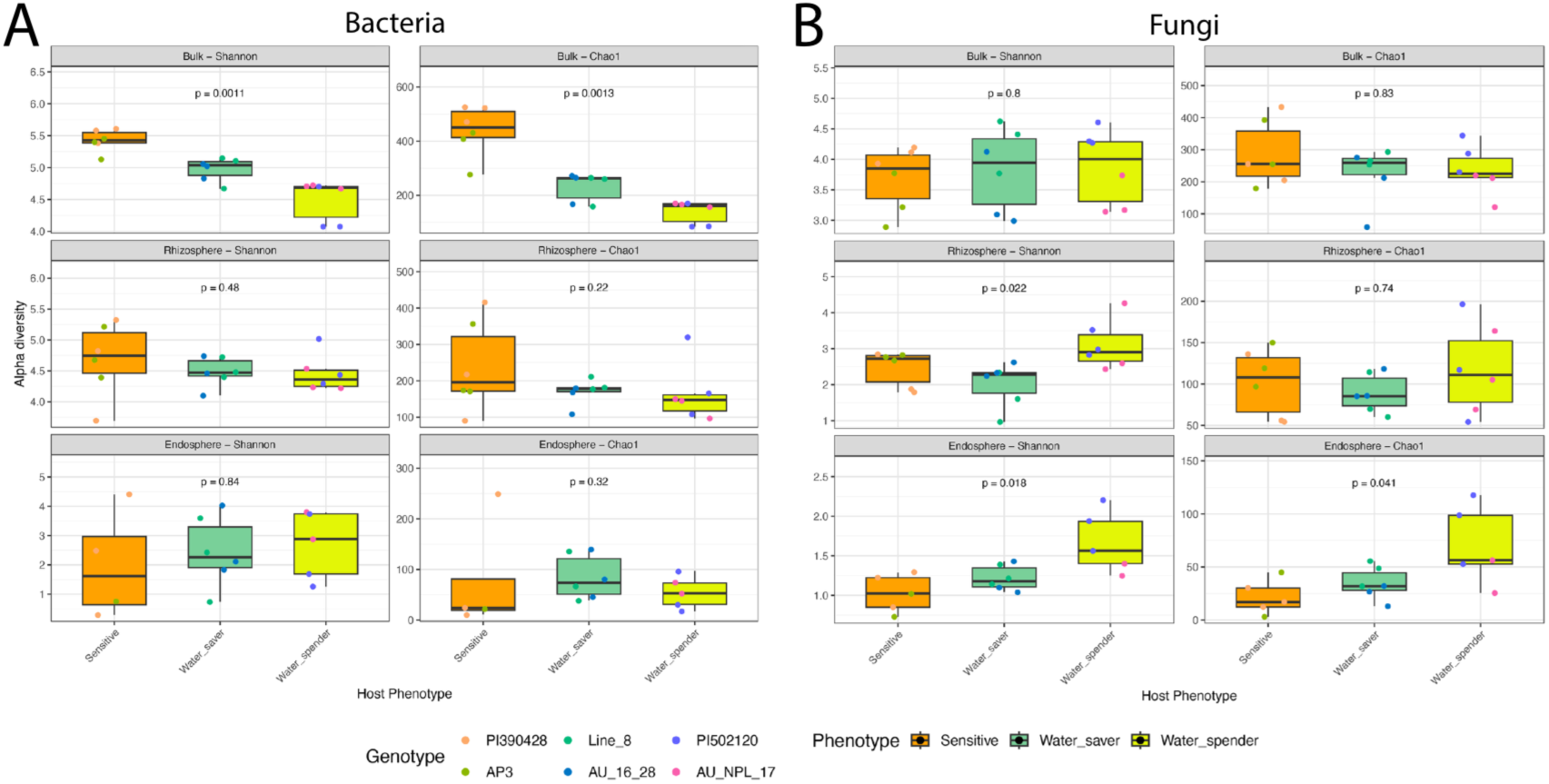
Alpha diversity of bacterial (16S) and fungal (ITS) communities across host phenotype groups for water-saver, water-spender, and sensitive peanut phenotypes. (A) Shannon Diversity (left) and Chao1 Richness (right) boxplots for bacterial communities in bulk soil (top) rhizosphere (middle) and root endosphere (bottom) plant compartments. (B) Shannon Diversity (left) and Chao1 Richness (right) boxplots for fungal communities in bulk soil (top) rhizosphere (middle) and root endosphere (bottom) plant compartments. Boxplots show the median, interquartile range (IQR) and whiskers extending to 1.5xIQR. Points beyond these whiskers are representative of outliers. Each host phenotype boxplot is a combination of samples from each host genotype of the phenotype grouping (n=3 per genotype, n=6 per phenotype, n=18 per plant compartment). Phenotype groupings within each plant compartment and diversity index were compared with Kruskal–Wallis tests followed by Dunn’s post hoc with a Benjamini–Hochberg correction with a significance threshold of (p < 0.05).

Patterns in the rhizosphere differed from those observed in bulk soil. Bacterial alpha diversity remained relatively stable across host groups (Table S2.1, Table S3.1), whereas fungal Shannon diversity varied significantly among both host phenotypes (KW, χ² = 7.614, df = 2, p = 0.022, Fig. 1; Table S5.1) and genotypes (KW, χ² = 11.246, df = 5, p = 0.047, Table S5.1). Pairwise comparisons indicated that the largest differences occurred between water-saver and water-spender phenotypes (p = 0.02; Table S5.2), indicating shifts in fungal community richness and evenness.

In the root endosphere, bacterial alpha diversity remained largely unchanged similar to observations in the rhizosphere. However, fungal communities exhibited significant differences in Shannon diversity among host phenotypes (KW, χ² = 6.899, df = 2, p = 0.018, Fig. 1; Table S6.1). These differences were driven primarily by contrasts between water-spender and drought-sensitive phenotypes (p = 0.03, Table S6.2).

### Evidence of Host Phenotype and Genotype effects on bulk soil Bacterial and Fungal Community composition

Bulk soil bacterial communities differed significantly among both host phenotypes and genotypes under Bray-Curtis and unweighted UniFrac distance metrics (Fig. S1 A-D). PERMANOVA analyses indicated that both factors explained significant variation in community composition, although genotype accounted for a larger proportion of variation than phenotype (PERMANOVA (16S Bulk Soil Bray-Curtis Host Phenotype): R² = 0.184, p = 0.0091; Host Genotype: R² = 0.375, p = 0.011, Fig. S1 A-B, Table S7.1). Despite overlap among groups in PCoA ordinations, CAP analyses showed clear separation among drought-response categories and genotypes (Fig 2A), with the strongest differentiation observed between water-saver and drought-sensitive hosts. Collectively, these results indicate that both drought-response phenotype and host genetic background are associated with bacterial community composition in bulk soil, with genotype exerting stronger influence.

**Figure 2:**
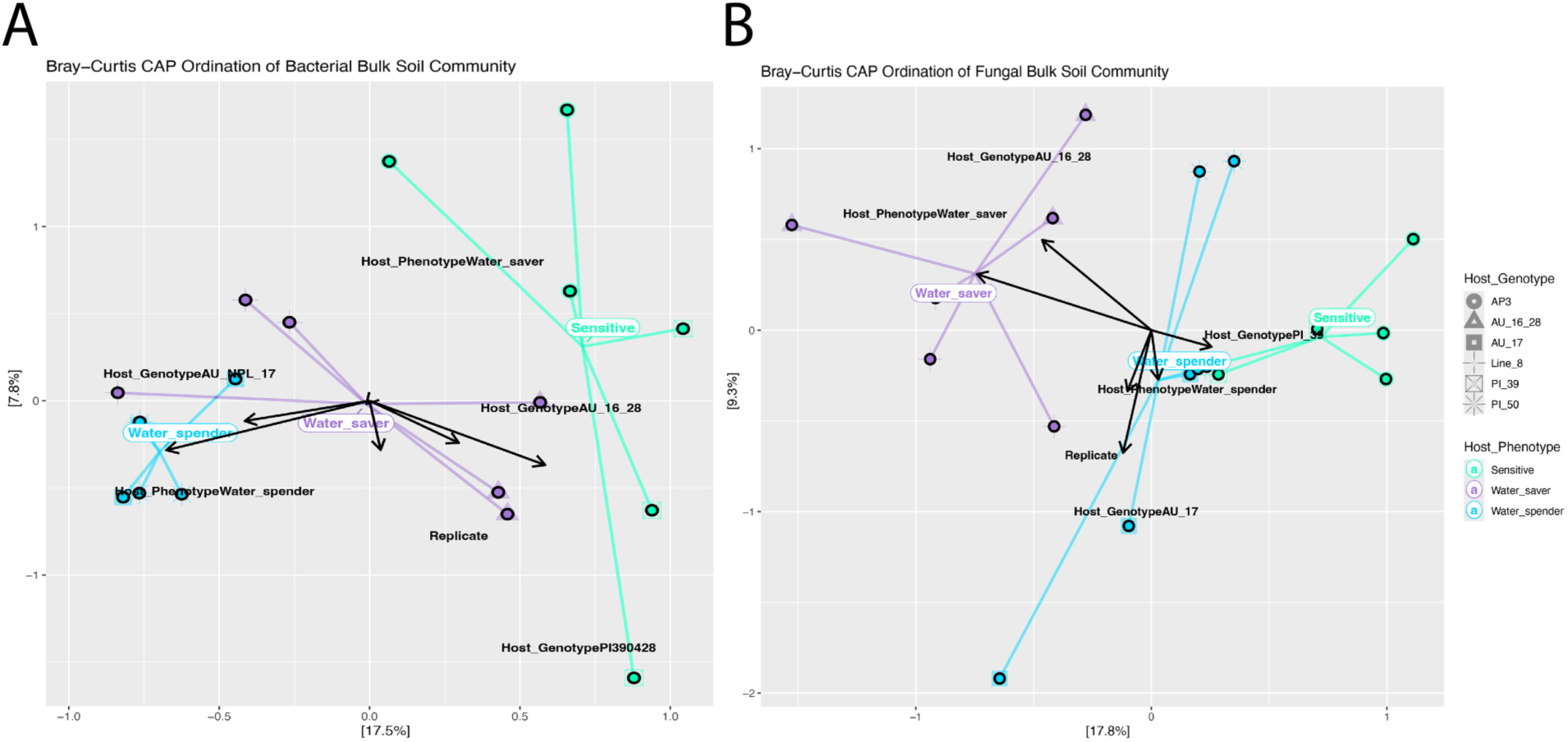
Canonical Analysis of Principal coordinates (CAP) plot of Bray-Curtis dissimilarities in bulk soil among (A) bacterial communities and (B) fungal communities. Host phenotype groupings are color coordinated and spider lines connect each sample of the host phenotype to the grouping centroid. Each dot represents one sample of each genotype (n=18 total, n=6 per phenotype). Percent variation that is explained by each axis is shown in brackets on the axis labels. Differences between host phenotype and host genotype in each plant compartment were evaluated using permutational multivariate analysis of variance (PERMANOVA), 9999 permutations). Genotype names in the CAP plot were abbreviated for visual clarity.

Similar patterns were observed for bulk soil fungal communities. Both host phenotype and genotype were associated with significant differences in fungal community composition under Bray-Curtis (Host Phenotype: R² = 0.2, p = 0.002, Host Genotype: R² = 0.42, p = <0.001, Table S8.1) and unweighted UniFrac (Host Phenotype: R² = 0.21, p = 0.023, Host Genotype: R² = 0.45, p = 0.006, Table S8.1) metrics. PCoA (Fig S2) and CAP (Fig 2B) ordinations revealed clustering by both phenotype and genotype, although genotype explained a larger proportion of community variation than phenotype. The consistency of these patterns across Bray-Curtis and UniFrac distances suggests that differences among host groups reflect shifts in both taxon abundances and the distribution of phylogenetically distinct fungal lineages. CAP ordinations further revealed directional clustering of samples according to drought-response category, with vectors separating water-saver, spender and drought-sensitive phenotypes despite substantial genotype-level variation, indicating a reproducible phenotype-associated component of community composition.

Alt text: Two side-by-side Bray-Curtis Canonical Analysis of Principal coordinates (CAP) plots visualizing the differences in phenotype grouping centroids of bacterial (16S) and fungal (ITS) communities.

Across both bacterial and fungal communities, significant phenotype-associated variation remained detectable despite substantial genotype-level effects. Together, these findings demonstrate that host identity, including both drought-response phenotype and genetic background, contributes to the assembly of bulk soil microbial communities. The generally stronger separation observed with Bray-Curtis distances further suggests that differences in taxon abundance were a major component of the observed community shifts (Fig.S1). Host genotype explains more variation in bulk soil communities, but a phenotype-associated signal remains detectable across multiple genetic backgrounds.

To distinguish the effects of drought-response phenotype from host genetic background, a nested PERMANOVA was performed with genotype nested within phenotype. Host phenotype explained a significant proportion of variation in bulk soil bacterial community composition (R^2^=0.184, p=0.002; Table S9.1), whereas genotype nested within phenotype was not significant (R^2^=0.191, p=0.130; Table S9.1) i.e. within individual phenotypic category, two genotypes do not differ significantly from one another. Host phenotype explained a significant proportion of variation in bulk soil fungal community composition (R^2^=0.205, p=0.001; Table S9.4), whereas genotype nested within phenotype was not significant (R^2^=0.217, p=0.017; Table S9.4) i.e. within individual phenotypic category, two genotypes differed significantly from one another. However, because only two genotypes represented each phenotype category, the power to detect genotype-within-phenotype effects was limited. To control for pseudoreplication effects that might contribute to this significant host phenotype effect, we ran a restricted model to block permutation recombination to host genotype. Results showed that the effect was no longer significant. To accurately represent the genotype replicates we wanted to test, we ran a genotype centroid model to average the technical replicates across genotype groupings. Results showed no significant model effect but there was a high reported level of variance associated with this result (R^2^ = 0.496, p= 0.2, Table S9.1 for bacteria and R^2^ = 0.452, p= 0.333, Table S9.4). These results indicate that differences among drought-response categories were more consistently associated with community composition than differences among genotypes within a given phenotype category.

### Host Phenotype and Genotype Effects persist in Rhizosphere and Root Endosphere fungal communities, but are weaker relative to bulk soil

In contrast to the bulk soil compartment, bacterial community composition in the rhizosphere and root endosphere was not significantly associated with either host phenotype or genotype under Bray-Curtis or unweighted UniFrac distance metrics (Fig. S1, Table 7.2, Table 7.3).

Although some analyses yielded moderate effect sizes, none were statistically significant, indicating weaker host-associated structuring of bacterial communities in these compartments. Fungal communities in the rhizosphere and root endosphere exhibited host-associated patterns, stronger than bacterial communities, but less pronounced compared to the fungal communities in the bulk soil compartment. In the rhizosphere, host genotype was significantly associated with fungal community composition under Bray–Curtis distances (PERMANOVA, R^2^ = 0.45, p = 0.028) (Table S8.2), whereas phenotype explained a moderate proportion of variation (R² = 0.20, Table S8.2), but did not reach statistical significance (p = 0.053, Table S8.2). Neither phenotype nor genotype significantly influenced fungal community composition under unweighted UniFrac distances.

In the root endosphere, fungal community composition differed significantly among host phenotypes under Bray-Curtis distances (R^2^ = 0.2755, p = 0.037, Table S8.3), whereas genotype effects were not significant. No significant differences were detected using unweighted UniFrac distances. Non-significant betadisper results across analyses indicated that observed differences were not driven by variation in group dispersion. Collectively, these findings suggest that host-associated effects on fungal communities are substantially reduced in the rhizosphere and root endosphere relative to bulk soil. Where significant differences were detected, they were observed primarily with Bray-Curtis distances, indicating that host-associated variation was largely driven by shifts in taxon abundance rather than turnover of phylogenetically distant fungal lineages.

### Relative Abundance patterns reveal phenotype-associated shifts in top 15 most abundant taxa

Alt text: Stacked multi-panel bar plot comparing the top 15 most abundant genera in each host phenotype grouping. There are two columns representing bacterial (16S) and fungal (ITS) phenotype plots that were separated into the three plant compartments: bulk soil, rhizosphere, and root endosphere.

The bacterial bulk soil community was dominated by *Sphingomonas, MND1*, and *Candidatus* Udaeobacter, which together accounted for the majority of sequences and remained relatively consistent across drought-response phenotypes (Fig 3A). Despite this overall stability, several lower-abundance taxa exhibited phenotype-associated patterns. Subgroup 10 was enriched in bulk soil compartment of drought-sensitive genotypes but was rare or absent in water-saver and water-spender phenotypes. *Acidibacter* was most abundant in water-spender soils. These shifts among lower-abundance taxa are consistent with the phenotype-associated differences detected in beta-diversity analyses. Although bacterial community composition in the rhizosphere and endosphere showed limited differences at the community level across phenotypes, several taxa exhibited lineage-specific abundance shifts. In the rhizosphere, dominant genera such as *Sphingomonas, Phenylobacterium, Burkholderiaceae*, *Novosphingobium* varied in relative abundance among drought-response phenotypes, whereas endosphere communities exhibited changes in relative abundance of some genera across phenotypes, with *Amycolatopsis* appearing enriched in water-saver and *Niastella*, *Dyella, Dokdonella,* and *Phenylobacterium* enriched in water-spender categories (Fig 3B, Fig 3C).

**Figure 3:**
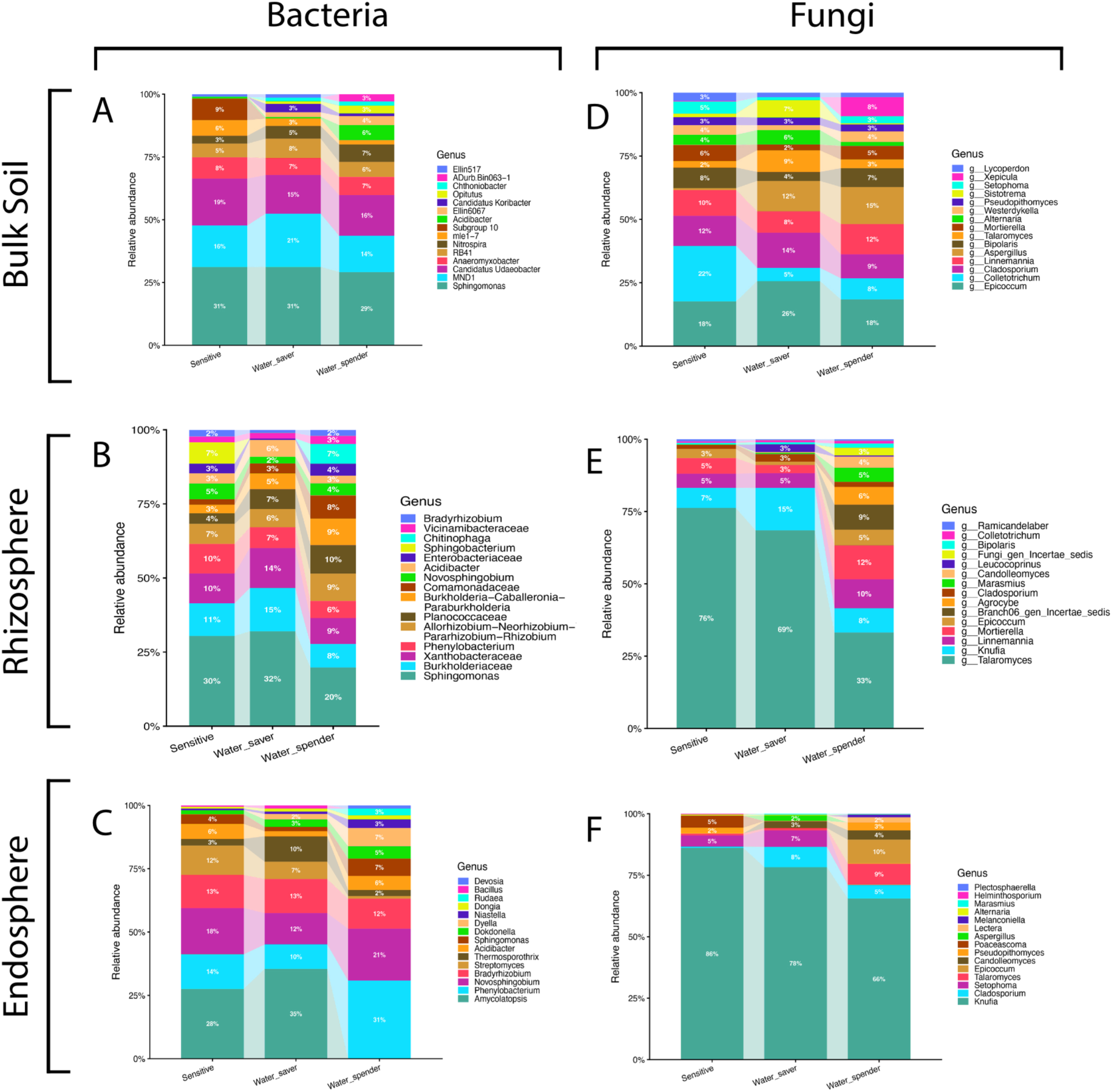
Relative abundance plots of the top 15 genera across host phenotypes categorized by (A) bacterial bulk soil (B) bacterial rhizosphere (C) bacterial root endosphere (D) fungal bulk soil (E) fungal rhizosphere and (F) fungal root endosphere. ASVs that were unclassified at the genus level were excluded prior to the plot generation. Bars show the mean relative abundance by each host phenotype for the respective compartments and kingdoms (n=6 per phenotype, n=18 per compartment), and abundances were renormalized so that the top 15 genera in each phenotype would sum to 100% of the bar; genera not represented in the top 15 are not included in the plots are thus did not contribute to the relative percentages found. Ribbons connecting each phenotype are strictly for visual comparison and do not represent any alterations to abundance percentages.

Bulk soil fungal communities displayed greater compositional variation among phenotypes. Although *Epicoccum*, *Colletotrichum*, and *Cladosporium* dominated all host groups, *Colletotrichum* showed substantial variation in abundance across phenotypes. In addition, *Sistotrema* was primarily associated with water-saver plants, whereas *Xepicula* was largely restricted to water-spender hosts (Fig 3D). Results for fungal rhizosphere abundance shifts reveal a more distinct pattern of host-associated community effects. While fungal communities were dominated by genus *Talaromyces* across all three phenotypes, its abundance varied markedly among host groups. Shifts in the next two most abundant genera observe a narrower spread in variance across phenotypes (*Knufia Linnemannia*. Several genera, including an unclassified *Branch 06 gen Incertae sedis, Agrocybe, Marasmius, and Candolleomyces* were enriched primarily in water-spender plants, consistent with the higher fungal diversity observed in this phenotype (Fig 3E). Similarly, fungal endosphere communities were dominated by *Knufia* across all phenotypes. However, water-spender hosts harbored a broader range of moderately abundant genera, including *Talaromyces, Epicoccum, Candolleomyces, Pseudopithomyces,* and *Lectera*, whereas *Poaceascoma* was detected primarily in drought-sensitive plants (Fig 3F).

These patterns are consistent with the elevated fungal diversity observed in water-spender rhizosphere and endosphere communities. Comparisons of fungal endosphere communities were interpreted cautiously due to unequal sample representation among phenotype groups.

### Differential Abundance Analysis Reveals Stronger Taxon-specific Responses in Fungal Communities

DESeq2 analyses revealed contrasting responses between bacterial and fungal communities. Despite significant phenotype-associated differences in bacterial bulk soil diversity and community composition, no bacterial taxa were consistently differentially abundant among phenotype groups. In contrast, multiple fungal taxa exhibited significant enrichment patterns, particularly in comparisons involving water-saver and drought-sensitive phenotypes. Consistent with relative abundance analyses, *Colletotrichum* was enriched in drought-sensitive plants relative to water-savers, whereas *Aspergillus* was enriched in both water-saver and water-spender phenotypes compared with drought-sensitive plants (Table 2).

**Table 2:** Differential Abundance Analysis Results for Fungal Communities by phenotype in bulk soil compartment.

Water-Saver vs Sensitive
| Phylum | Family | Genus | Log2 Fold Change | Adjusted p-value |
| --- | --- | --- | --- | --- |
| *Ascomycota* | *Glomerellaceae* | *Colletotrichum* | -2.31 | 0.0145 |
| *Ascomycota* | *Aspergillaceae* | *Aspergillus* | 4.55 | 0.0145 |
| *Ascomycota* | *Nectriaceae* | *Fusarium* | -6.67 | 0.0145 |
| *Ascomycota* | *Tubeufiaceae* | *Helicoma* | -4.79 | 0.0145 |
| *Ascomycota* | *Trichocomaceae* | *Talaromyces* | 2.05 | 0.0189 |
| *Chytridiomycota* | *Powellomycetaceae* | *Fimicolochytrium* | -3.80 | 0.0189 |
| *Ascomycota* | *Phaeosphaeriaceae* | *Phaeosphaeria* | -2.86 | 0.0190 |

Water-Spender vs Sensitive
| Phylum | Family | Genus | Log2 Fold Change | Adjusted p-value |
| --- | --- | --- | --- | --- |
| *Ascomycota* | *Aspergillaceae* | *Aspergillus* | 5.13 | 0.00534 |

Water-Saver vs Water-Spender
| Phylum | Family | Genus | Log2 Fold Change | Adjusted p-value |
| --- | --- | --- | --- | --- |
| *Ascomycota* | *Tubeufiaceae* | *Helicoma* | -5.40 | 0.00785 |
| *Ascomycota* | *Nectriaceae* | *Fusarium* | -6.43 | 0.01570 |

Several additional differentially abundant taxa were detected despite being absent from the top 15 most abundant genera, indicating that phenotype-associated shifts extended beyond dominant community members. Many of these changes occurred within members of the phylum *Ascomycota* (Table 2).

In the rhizosphere, no bacterial taxa were differentially enriched among phenotypes. In contrast, the fungal genus *Agrocybe* was strongly enriched in water-spender relative to water-saver genotypes (Table 3). Within the endosphere, the pattern was reversed. No fungal taxa were significantly enriched, whereas the bacterial genus *Amycolatopsis* was enriched in water-saver relative to water-spender hosts (Table 3). Together, these findings suggest that while broad community-level differences were most evident in bulk soil, host-associated effects in rhizosphere and endosphere compartments were largely restricted to a small number of responsive microbial lineages.

**Table 3:** Differential Abundance Analysis Results of fungal and bacterial taxa between water-saver and -spender in rhizosphere and endosphere.

Water-Saver vs Water-Spender: Fungal Rhizosphere
| Phylum | Family | Genus | Log2 Fold Change | Adjusted p-value |
| --- | --- | --- | --- | --- |
| *Basidiomycota* | *Strophariaceae* | *Agrocybe* | -8.88 | 0.0107 |

Water-Saver vs Water-Spender: Bacterial Endosphere
| Phylum | Family | Genus | Log2 Fold Change | Adjusted p-value |
| --- | --- | --- | --- | --- |
| *Actinobacteriota* | *Pseudonocardiaceae* | *Amycolatopsis* | 9 | 0.000464 |

### Sloan neutral community model analysis predicts strong niche differentiation in fungal community assembly between phenotypes

Sloan’s neutral fit model revealed marked differences in assembly processes between bacterial and fungal communities across plant compartments. Bacterial communities were generally well described by the neutral model, particularly in bulk soil and rhizosphere compartments where model fit was high, with bulk soil (R^2^ = 0.875 – 0.929) and rhizosphere (R^2^ = 0.753 – 0.936) exhibiting high R^2^ ranges (Table 4). In contrast, fungal communities in bulk soil (R^2^ = 0.37 – 0.5), rhizosphere (R^2^ = -0.003 – 0.387), and endosphere (R^2^ = -0.398-0.463) consistently exhibited lower model fit, indicating a greater influence of deterministic processes on fungal communities (Table 4). Consistent with this pattern, fungal communities contained substantially more taxa that deviated from neutral expectations than bacterial communities, suggesting that fungal assembly was less governed by stochastic dispersal and drift (Table S10; Table S11).

**Table 4:**
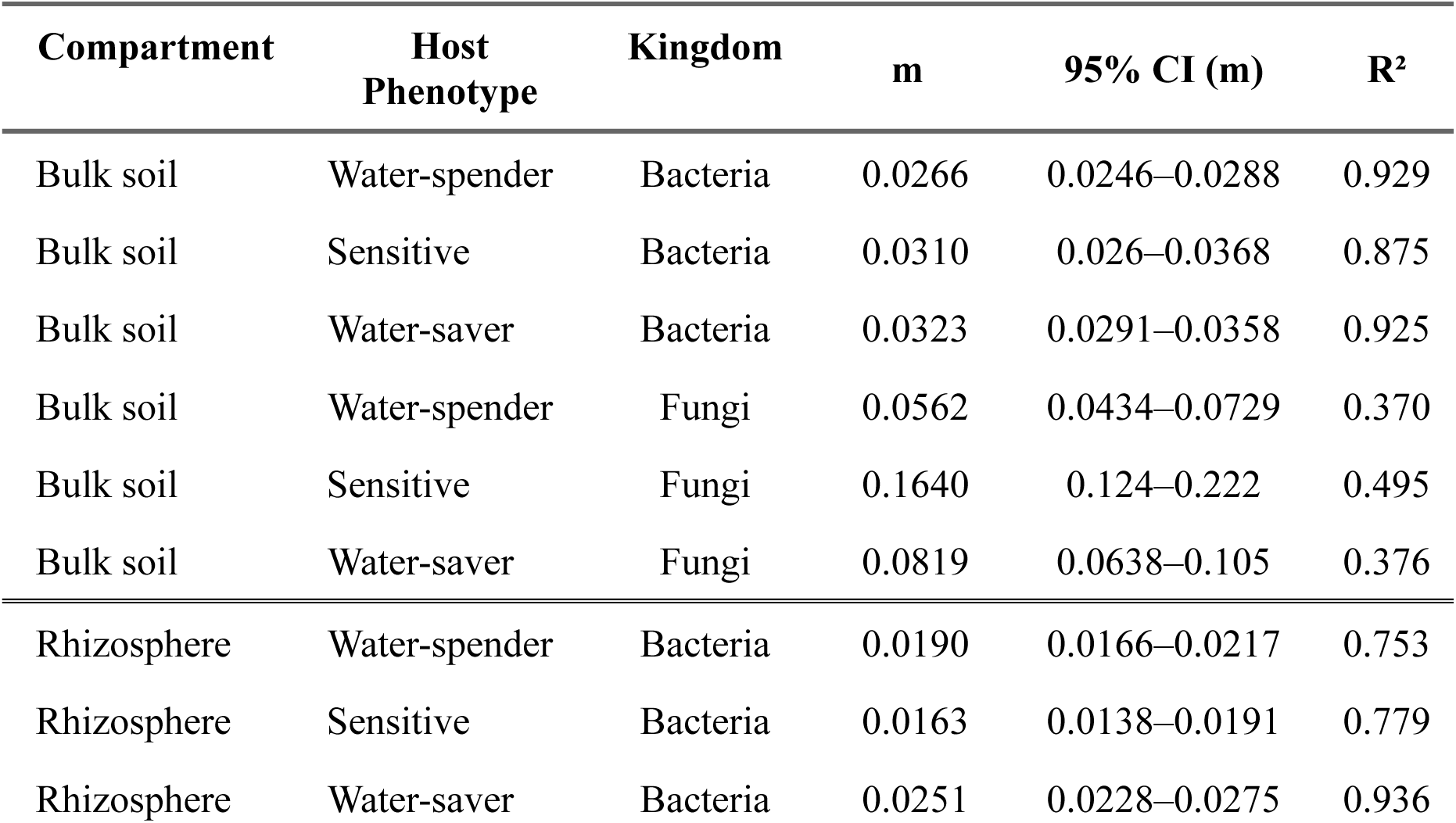
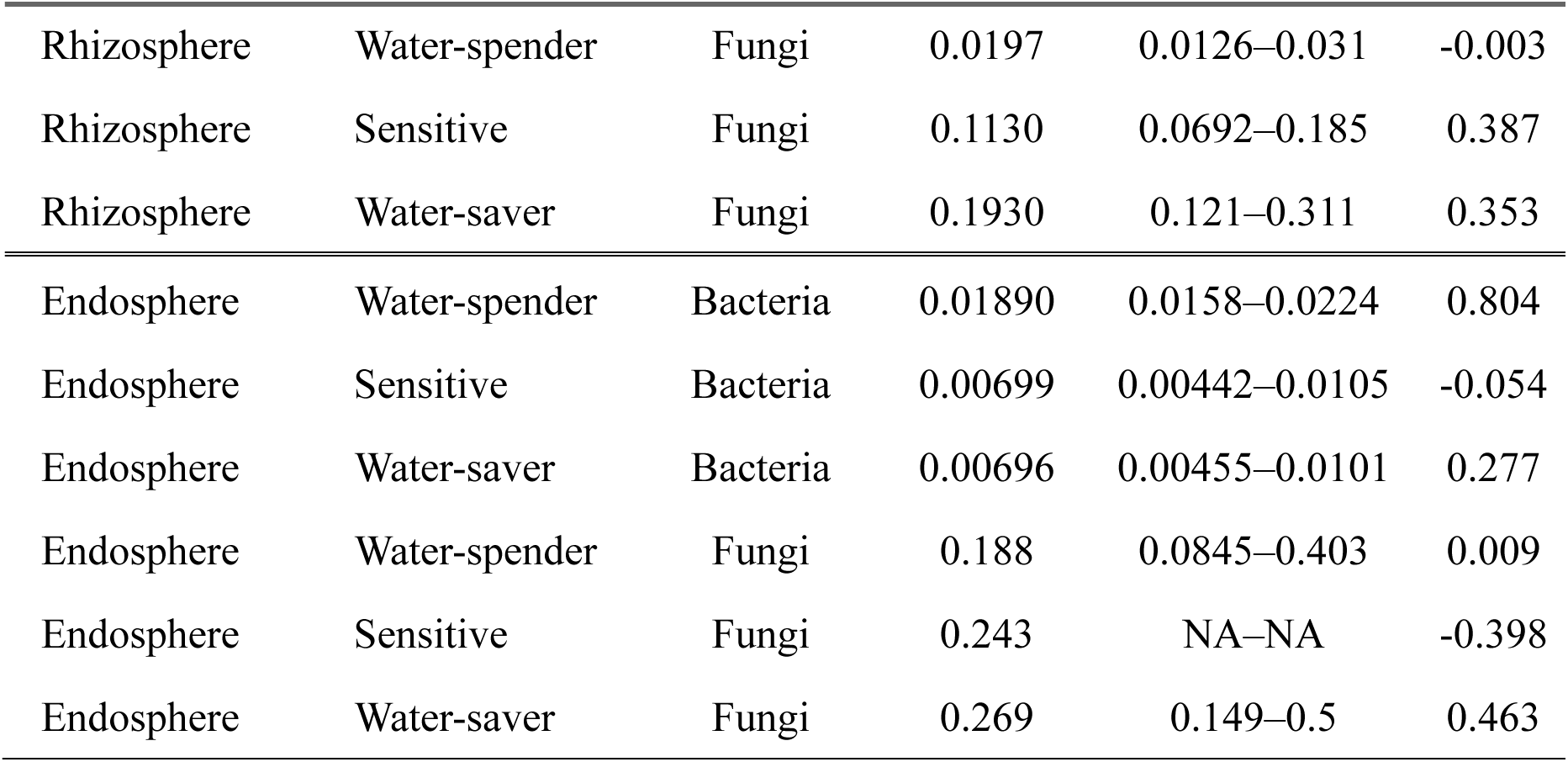
Sloan neutral community model fit statistics of bacterial/fungal communities in each plant compartment by host Phenotype. m, migration rate; R^2^, proportion of variance in occurrence frequency explained by the neutral model.

The relative importance of neutral processes also varied by plant compartments. For bacterial communities, neutral model fit was highest in bulk soil (R^2^ = 0.875 – 0.929) and remained high in the rhizosphere (R^2^ = 0.753 – 0.936) indicating that stochastic processes contributed substantially to community assembly in soil-associated habitats. However, model fit of bacterial communities declined in the endosphere of sensitive and water-saver genotypes (R^2^ = -0.054 and 0.277, respectively; Table 4), suggesting stronger host filtering within root tissues. In contrast, fungal communities showed weak support for neutral assembly across all compartments, with rhizosphere and endosphere communities, exhibiting near-zero and negative R^2^ values (Table 4). Neutral processes alone could not explain observed taxon abundance patterns.

Across all plant compartments, deterministic factors (factors outside of random dispersal/drift predictions based on abundance) acted on fungal communities more than bacterial communities. Significantly, fungal bulk soil had the highest number of suppressed taxa across all phenotypes (below-prediction 39/above-prediction taxa: 8) and fungal rhizosphere communities had the highest number of enriched taxa across all phenotypes (below-prediction taxa: 27/above-prediction taxa: 19), showing a clear pattern of deterministic assembly occurring differentially between microbial kingdoms (Fig. 4). All three phenotypes had large communities of suppressed fungal taxa in their model fits, with water-spender suppressing bulk soil genera such as *Agrocybe sp,, Aspergillis sp.,* and surprisingly, *Mortierella sp* (Table S11). Enriched bulk soil taxa in water-spenders included *Rhizopus arrhizus, Torula sp.,* and *Spizellomycetales_gen_Incertae_sedis sp.* (Table S11). The highest level of fungal enrichment in the rhizosphere was in association with water-spender phenotypes; taxa included but not limited to *Allophoma tropica, Candolleomyces luteopallidus,* and several species *of Colletotrichum* were modelled to be uniquely enriched in water-spender (Table S11). In the rhizosphere, water-saver exhibited lower levels of enriched taxa that fit Sloan’s neutral community model, with *Hannaella oryzae* being the only species (Table S11). All bacterial taxa fit the neutral model in bulk soil, with one taxa (*Chitinophaga sp.*) being suppressed in the rhizosphere of the water-spender phenotype grouping (Table S10). In the endosphere, similar patterns of highly selective enrichment/suppression were seen across bacterial and fungal communities, with the exception of water-spender phenotypes that exhibited higher levels of fungal taxa suppression (*Epicoccum pimpprinum, Lectera nordwiniana,* and unclassified genera from the *phyla Ascomycota)* (Table S11). Together, these findings suggest that host phenotype imparts selective recruitment processed that might favor shifts in fungal taxa, particularly in bulk and rhizosphere soil compartments.

**Figure 4:**
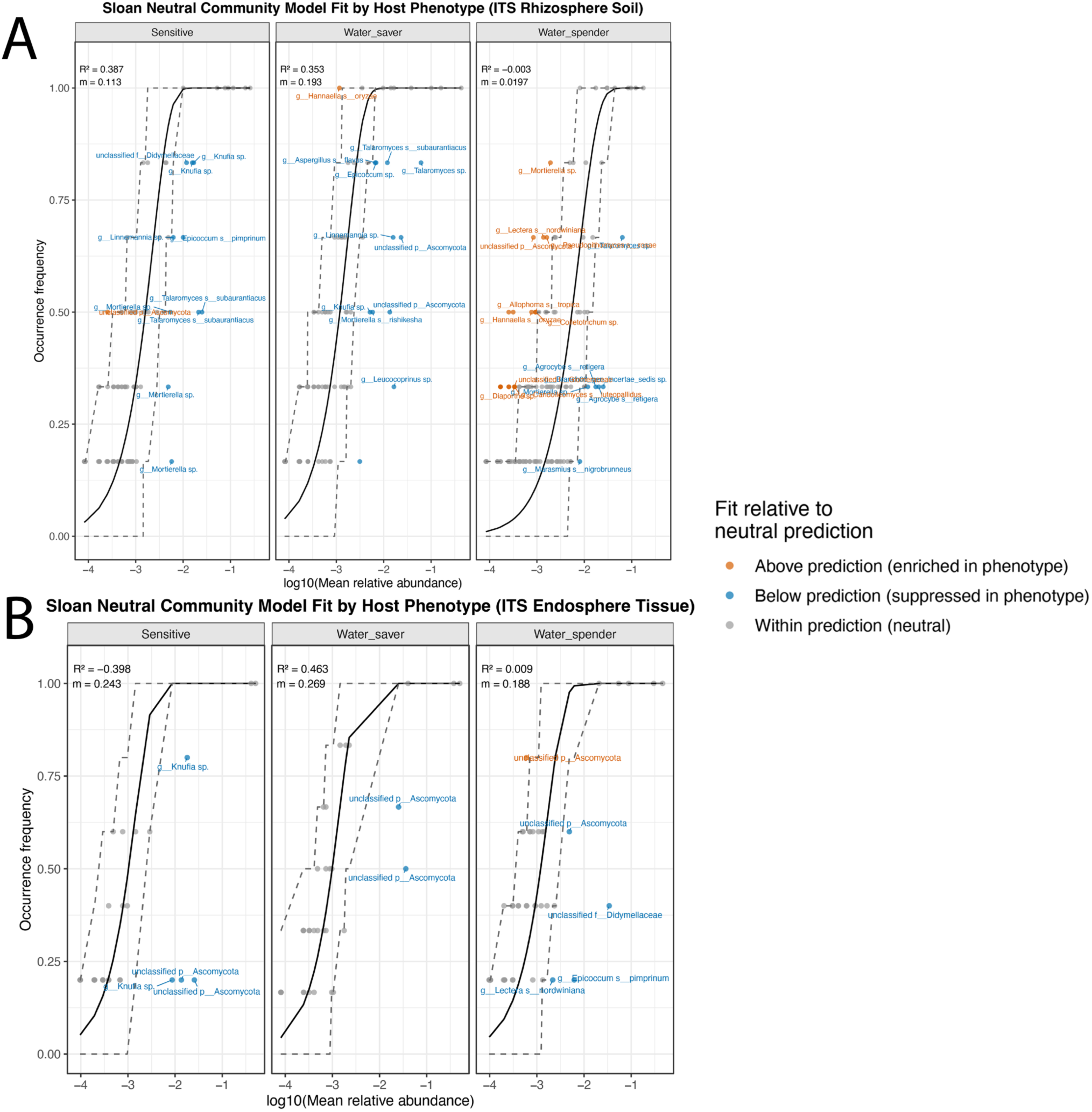
Fit of the Sloan neutral community model for (A) fungal rhizosphere communities and (B) fungal root endosphere communities separated by host phenotype. Each point on the plot represents one ASV plotted by mean relative abundance (log10 scale, x-axis). The solid black line represents the best fit to the community model, with the dashed lines representing the 95% confidence interval of best fit predictions for each respective phenotype grouping. ASVs that occur less frequently in the phenotype grouping than what is predicted by abundance data alone is represented in blue, and ASVs that occur more frequently are shown in orange. R^2^ values indicated the goodness of fit, with a lower value indicating community assembly being highly influenced by niche differentiation/non-stochastic processes. M values represent migration rate, with a lower m value indicating limited dispersal predictions.

Host phenotype also influenced the balance between neutral and deterministic assembly within each plant compartment. Among bacterial rhizosphere communities, water-saver genotypes exhibited the highest neutral model fit (R^2^=0.936; Table 4). In contrast, fungal communities associated with water-spender genotypes showed little to no neutral signal in the rhizosphere (R^2^ near zero), indicating that deterministic processes dominated fungal recruitment in these plants. There was a progressive shift in stochastic assembly in bulk soil towards increasingly deterministic assembly in root-associated compartments (Table 4) with this transition being stronger for fungi than for bacteria (Table 4).

Alt text: Two scatter plot panels showing the Sloan neutral community model for (A) fungal rhizosphere communities and (B) fungal endosphere communities separated by host phenotype.

## Discussion

### Drought-response phenotype is associated with reproducible microbiome variation across peanut genotypes

Host genotype is a recognized determinant of plant microbiome assembly, although its effects are intertwined with environmental conditions and host functional traits (Bulgarelli et al. 2013; Walters et al. 2018; Bergelson et al. 2019). In this study, we sought to determine whether peanut genotypes exhibiting contrasting drought-response strategies harbor distinct microbial communities and whether these patterns are detectable across multiple genetic backgrounds.

Consistent with our hypothesis, both drought-response phenotype and host genotype were associated with variation in microbial community structure, particularly within the bulk soil compartment. Importantly, phenotype-associated patterns remained detectable despite underlying genetic variation among genotypes, suggesting that host traits linked to drought response contribute to microbiome assembly beyond individual genotype effects. Several studies have demonstrated that plant-associated microbiomes are influenced not only by host taxonomic identity but also by host functional traits (Fitzpatrick et al. 2018; Trivedi et al. 2020; Deng et al. 2021). Our findings extend these observations to peanut by showing that microbial community differences were associated with water-saver, water-spender, and drought-sensitive host categories under non-stress conditions. Our findings suggest that drought-response phenotypes act as distinct ecological niches capable of supporting different microbial assemblages. Notably, these associations were detected under field conditions, and across multiple genetic backgrounds, indicating that host traits linked to drought adaptation contribute to microbiome assembly under agronomically realistic conditions.

### Phenotype-associated signals persist despite host genetic background effects

A central challenge in plant microbiome studies is disentangling host phenotype effects from underlying genetic background. To address this issue, we included two genotypes within each drought-response category and explicitly evaluated genotype and phenotype effects. Separate PERMANOVA analyses indicated that both factors significantly influenced bulk soil bacterial and fungal community composition, with genotype often explaining a larger proportion of variation. However, nested PERMANOVA provided additional insight. The nested PERMANOVA revealed a significant phenotype effect but no significant genotype-within-phenotype effect, suggesting that bacterial community differences observed in bulk soil were associated with drought-response categories across genetic backgrounds. While the limited number of genotypes representing each phenotype category reduced the power to detect genotype-within-phenotype effects, these results indicate that microbial community differences were more consistently associated with drought-response categories. Similar observations have been reported in maize, rice, *Arabidopsis*, and sorghum, where shared host functional characteristics can shape microbial community assembly across genetically distinct accessions (Edwards et al. 2015; Walters et al. 2018; Wagner et al. 2016; Deng et al. 2021). While additional genotypes would be needed to fully disentangle phenotype and genetic background effects, these results support the existence of phenotype-associated component of microbiome assembly.

### Root exudation and host functional traits may mediate phenotype-associated microbiome assembly

The mechanisms underlying these phenotype-associated microbial patterns remain unclear. However, differences in root exudation and root system architecture provide a plausible explanation. Root exudates are among the primary determinants of rhizosphere microbial assembly because they supply carbon substrates and signaling compounds that regulate microbial recruitment, competition, and persistence (Badri and Vivanco, 2009; Sasse et al. 2018; Jacoby et al. 2020). Water-saver and water-spender genotypes examined in this study differ in physiological characteristics associated with drought adaptation, including stomatal regulation, water-use patterns, and root system development. Such differences could influence both the quantity and composition of carbon released into the surrounding soil, creating distinct microbial niches. Specifically, deeper root system in water-spender is a heritable trait observed regardless of drought stress and such root phenotype may generate larger rhizosphere effect and greater carbon inputs, supporting more diverse microbial communities. Consistent with this possibility, water-spender varieties exhibited greater fungal diversity and contained several uniquely enriched fungal genera in rhizosphere and endosphere. Although exudates were not measured in this study, future integration of metabolomics and microbiome profiling can help address whether drought-response phenotypes generate distinct exudates that shape microbial recruitment.

### Bulk soil exhibited the strongest phenotype-associated microbial patterns

One of the more unexpected findings was that the strongest phenotype- and genotype-associated microbial patterns occurred in the bulk soil compartment. Classical models of plant microbiome assembly predict stronger host effects in the rhizosphere and endosphere habitats because these compartments experience increasing levels of host filtering (Bulgarelli et al. 2013; Compant et al. 2019). However, both bacterial and fungal communities exhibited their strongest phenotype-and genotype-associated shifts in bulk soil, whereas host-associated effects were generally weaker in rhizosphere and endosphere communities.

Bulk soil functions as a highly diverse “reservoir” of recruitable microbes for the rhizosphere and root endosphere compartments. Consequently, shifts within this compartment can influence downstream colonization process closer to the root. A potential explanation for the strong signature observed in the bulk soil is that long-term interactions between plants and soil generate plant-mediated legacy effects that extend beyond the immediate root zone. Differences in root growth and rhizodeposition can progressively alter bulk soil microbial communities over time (de Vries et al. 2020). The stronger phenotype-associates signal in bulk soil may therefore reflect cumulative effects of contrasting drought-response strategies on soil microbial habitats rather than direct root selection alone. Furthermore, Bray-Curtis distances generally produced stronger phenotypic group separation than unweighted UniFrac distances across both bacterial and fungal communities. This pattern suggests that differences among drought-response categories were driven primarily by shifts in relative abundance of existing taxa rather than wholesale turnover of phylogenetically distinct lineages. Such abundance-driven filtering is consistent with ecological filtering mechanisms that alter the success of specific microbial groups, in order to cater specifically to what the phenotype in question might need for success (i.e the potential for greater nutrient acquisition from water-spender phenotypes), without fundamentally changing overall community membership. The alternative would be acquiring entirely new dominant taxa specific to each phenotype, which was not supported by this study. This conclusion was further supported by DESeq2 results which showed a surprising lack of differentially abundant microbes in bacterial bulk soil communities, despite their role in being highly adaptable to changes in nutrient filtering processes.

### Fungal communities were more responsive than bacterial communities within plant-associated compartments

Fungal communities exhibited stronger phenotype-associated responses than bacterial communities across plant-associated compartments. While beta-diversity analyses indicated that fungal communities were primarily differentiated by shifts in the relative abundance of closely related taxa rather than wholesale turnover of phylogenetically distinct lineages, differential abundance analyses revealed clear phenotype-specific enrichment patterns. Several of the most differentially abundant fungal taxa were also among the dominant genera within bulk soil communities, indicating that drought-response phenotypes influenced both overall community structure and the abundance of ecologically important taxa. Consistent with these observations, Sloan neutral community modeling indicated stronger deterministic assembly of fungal communities than bacterial communities across plant compartments.

Several fungal taxa displayed phenotype-specific enrichment patterns that may reflect differences in host traits associated with drought adaptation. For example, *Colletotrichum*, *Rhizophydiales, Sporobolomyces* were predicted to occur above neutral expectations only in water-spender phenotypes. The deeper and more extensive root systems may create conditions that favor recruitment of specific fungal lineages. Similarly, *Agrocybe* was enriched in water-spender rhizosphere communities and has been associated with improved nutrient acquisition and plant growth in other systems. This species has been studied as an inoculant to aid in plant growth parameters and total P and N uptake from the substrate by indirectly aiding in nutrient acquisition processes (Vohník et al., 2012). Associations between root length and deterministic assembly of fungal taxa have further been studied in temperature grassland species, and in this study, we hypothesize that these specific drought-tolerant root characteristics directly influenced the assembly of unique fungal taxa that could aid in critical processes of nutrient acquisition and drought resistance (Sweeney et al., 2021).

Previous studies in other host systems have credited host phenotype with altering the rhizosphere bacterial communities due to their deterministic nature within the plant compartment (Hodgson et al., 2025). In this study, we observed only a few bacterial taxa to be specifically associated with water-saver or water-spender phenotypes. For example, *Amycolatopsis* enriched in the endosphere of water-saver peanut genotypes, or *Dyella* enriched in endosphere of water-spender genotypes. The stronger response of fungal communities relative to bacterial communities in the rhizosphere and endosphere compartments suggests that fungal communities may be more sensitive to variation in root-derived resources and other environmental filters. Notably, the magnitude of phenotype effects differed depending on the level of analysis (DESeq2, and beta diversity analysis). While community-level differences in rhizosphere and endosphere composition were generally weaker than those observed in bulk soil, differential abundance analyses identified specific microbial taxa that were enriched within particular drought-response phenotypes. This pattern suggests that assembly processes in the rhizosphere and endosphere may be influenced not only by host-mediated selection but also by other factors such as edaphic conditions and may operate through targeted responses of individual microbial lineages rather than broad restructuring of entire communities (Bulgarelli et al. 2013; Edwards et al. 2015).

Although the mechanisms underlying the differences among microbial community assemblies across drought response phenotypes remain unclear, they likely reflect interactions among host-mediated selection, soil physicochemical properties, moisture availability, and nutrient status.

Given the low biomass and increased sparsity typical of endosphere datasets, such single-taxon signals should be interpreted cautiously as they may reflect both biological specificity and statistical sensitivity to rare taxa. Future studies integrating root exudate profiling, soil physicochemical measurements, metagenomics, and transcriptomics would shed light on the functional connection between phenotype influence and microbial processes that are selected for by the phenotype.

## Conclusion

This study demonstrates that peanut drought-response phenotypes are associated with reproducible patterns of bacterial and fungal community assembly across multiple host genetic backgrounds, even in the absence of drought stress under field conditions. Although host genotype contributed substantially to microbiome variation, phenotype-associated signal remained evident, particularly within bulk-soil communities and root-associated fungal assemblages. These findings suggest that plant functional traits linked to drought adaptations such as root architecture, water-use strategies, carbon allocation, and potentially root exudation patterns, shape microbiome assembly prior to the onset of stress. Stronger abundance-based responses relative to phylogenetic turnover further suggest that drought-response phenotypes primarily influence the relative abundance of existing microbial taxa rather than recruiting entirely distinct microbial lineages.

From a breeding perspective, these results highlight the potential for drought-adaptive phenotypes to influence plant physiology and composition of associated microbial communities. As efforts increasingly integrate host genetics and microbiome-based approaches to improve crop resilience, understanding the mechanisms underlying phenotype-associated microbiome assembly will become increasingly important. Future studies combining host genomics, root phenotyping, exudate metabolomics, and microbiome manipulation experiments will be necessary to determine whether the microbial signatures identified here are consequences of drought-adaptive traits or active contribution to drought resilience in peanut. Screening of differentially enriched fungal taxa for root exudate modulation and assessing the biological and chemical changes to the host environment will establish further insight into the role that these taxa might play in enhancing the overall drought-resistance mechanisms of specialized peanut genotypes. The findings from this work highlight the potential for integrating microbiome-informed approaches into future drought-resilience breeding programs.

## Data Availability

The raw reads from this project have been submitted to the SRA database under the BioProject accession number PRJNA1493213. All the codes used in the analysis presented in this study are available on the GitHub repository https://github.com/Potnislab/peanut_genotype_microbiome_manuscript

## Acknowledgements

This work was funded by the National Science Foundation under Grant Award No. (2316278). Any opinions, findings, and conclusions or recommendations expressed in this material are those of the author(s) and do not necessarily reflect the views of the National Science Foundation.

Special thanks to Dr. Chen and colleagues at the Peanut Breeding Program at Auburn University for providing plants to sample and assisting during sampling.

This study used Claude (Anthropic, Sonnet 4.6) and ChatGPT (OpenAI, GPT-5.5) to assist in debugging of R code used for phyloseq, vegan, ggplot2 and DESeq2 workflows. After using these tools, ZL carefully confirmed, reviewed, and edited the content of the code and takes full responsibility for the content of the publication.

## Author Contribution Statement

NP: conceptualization; ZJL, DS and NP: methodology; ZJL: formal analysis; ZJL, DS and NP: investigation; CC and NP: resources; ZJL: data curation; ZJL, and NP: writing - original draft; ZJL, CC, and NP: writing - review & editing; ZJL, and NP: visualization; NP: supervision; NP: funding acquisition

## Supplementary data

The following supplementary data are available at JXB online.

Fig S1: Principal coordinate analysis of all bacterial communities grouped by plant compartment with phenotype and genotype comparisons in each compartment.

Fig S2: Principal coordinate analysis of all bacterial communities grouped by plant compartment with phenotype and genotype comparisons in each compartment.

Fig. S3: Sampling design schematic.

Table S1.1: Alpha diversity Kruskal-Wallis results for bacterial communities in bulk soil.

Table S1.2: Dunn Post-hoc pairwise comparison results for bacterial bulk soil communities by phenotype.

Table S1.3: Dunn Post-hoc pairwise comparison results for bacterial bulk soil communities by genotype.

Table S2.1: Alpha diversity Kruskal-Wallis results for bacterial communities in rhizosphere soil.

Table 3.1: Alpha diversity Kruskal-Wallis results for bacterial communities in root endosphere.

Table 4.1: Alpha diversity Kruskal-Wallis results for fungal communities in bulk soil.

Table S4.2: Dunn Post-hoc pairwise comparison results for fungal bulk soil communities by genotype.

Table S5.1: Alpha diversity Kruskal-Wallis results for fungal communities in rhizosphere soil.

Table S5.2: Dunn Post-hoc pairwise comparison results for fungal rhizosphere soil communities by phenotype.

Table S5.3: Dunn Post-hoc pairwise comparison results for fungal rhizosphere soil communities by genotype.

Table S6.1: Alpha diversity Kruskal-Wallis results for fungal communities in root endosphere.

Table S6.2: Dunn Post-hoc pairwise comparison results for fungal root endosphere communities by phenotype.

Table S7.1: Bacterial (16S) Bulk Soil Beta Diversity Results for Bray-Curtis and Unweighted Unifrac distance metrics.

Table S7.2: Bacterial (16S) Rhizosphere Soil Beta Diversity Results for Bray-Curtis and Unweighted Unifrac distance metrics.

Table S7.3 Bacterial (16S) Root Endosphere Beta Diversity Results for Bray-Curtis and Unweighted Unifrac distance metrics.

Table S8.1: Fungal (ITS) Bulk Soil Beta Diversity Results for Bray-Curtis and Unweighted Unifrac distance metrics.

Table S8.2: Fungal (ITS) Rhizosphere Soil Beta Diversity Results for Bray-Curtis and Unweighted Unifrac distance metrics.

Table S8.3: Fungal (ITS) Root Endosphere Beta Diversity Results for Bray-Curtis and Unweighted Unifrac distance metrics.

Table S9.1. Nested PERMANOVA model results for bacterial (16S) bulk soil community.

Table S9.2. Nested PERMANOVA model results for bacterial (16S) rhizosphere soil community.

Table S9.3. Nested PERMANOVA model results for bacterial (16S) root endosphere community.

Table S9.4. Nested PERMANOVA model results for fungal (ITS) bulk soil community.

Table S9.5. Nested PERMANOVA model results for fungal (ITS) rhizosphere soil community.

Table S9.6. Nested PERMANOVA model results for fungal (ITS) root endosphere community.

Table S10. Sloan neutral community model fit statistics of bacterial (16S) bulk soil, rhizosphere, and root endosphere communities by host Phenotype.

Table S11. Sloan neutral community model fit statistics of fungal (ITS) bulk soil, rhizosphere, and root endosphere communities by host Phenotype.

