## Supplementary material for "Drought-tolerant phenotypes associated with patterns of deterministic microbiome assembly across peanut genotypes": Table S1, Table S2, Table S3, Table S4, Table S5, Table S6, Table S7, Table S8, Table S9, Table S10, Table S11, Fig S1, Fig S2, Fig S3

*Figure S1: PCoA plots of community dissimilarity analysis for bacterial communities for each plant compartment grouped by phenotype using Bray-Curtis (A, E, I) and Unweighted Unifrac (C, G, K); additional grouping by genotype was performed using Bray-Curtis (B, F, J) and Unweighted Unifrac (D, H, L) Phenotype analyses are grouped by centroid ellipses (n = 3 per genotype, n = 6 per phenotype). The x-axis represents PCoA1 that accounts for the largest variation and the y-axis represents PCoA2 accounting for the second largest variation.*

*Figure S2: PCoA plots of community dissimilarity analysis for bacterial communities for each plant compartment grouped by phenotype using Bray-Curtis (A, E, I) and Unweighted Unifrac (C, G, K); additional grouping by genotype was performed using Bray-Curtis (B, F, J) and Unweighted Unifrac (D, H, L) Phenotype analyses are grouped by centroid ellipses (n = 3 per genotype, n = 6 per phenotype). The x-axis represents PCoA1 that accounts for the largest variation and the y-axis represents PCoA2 accounting for the second largest variation.*

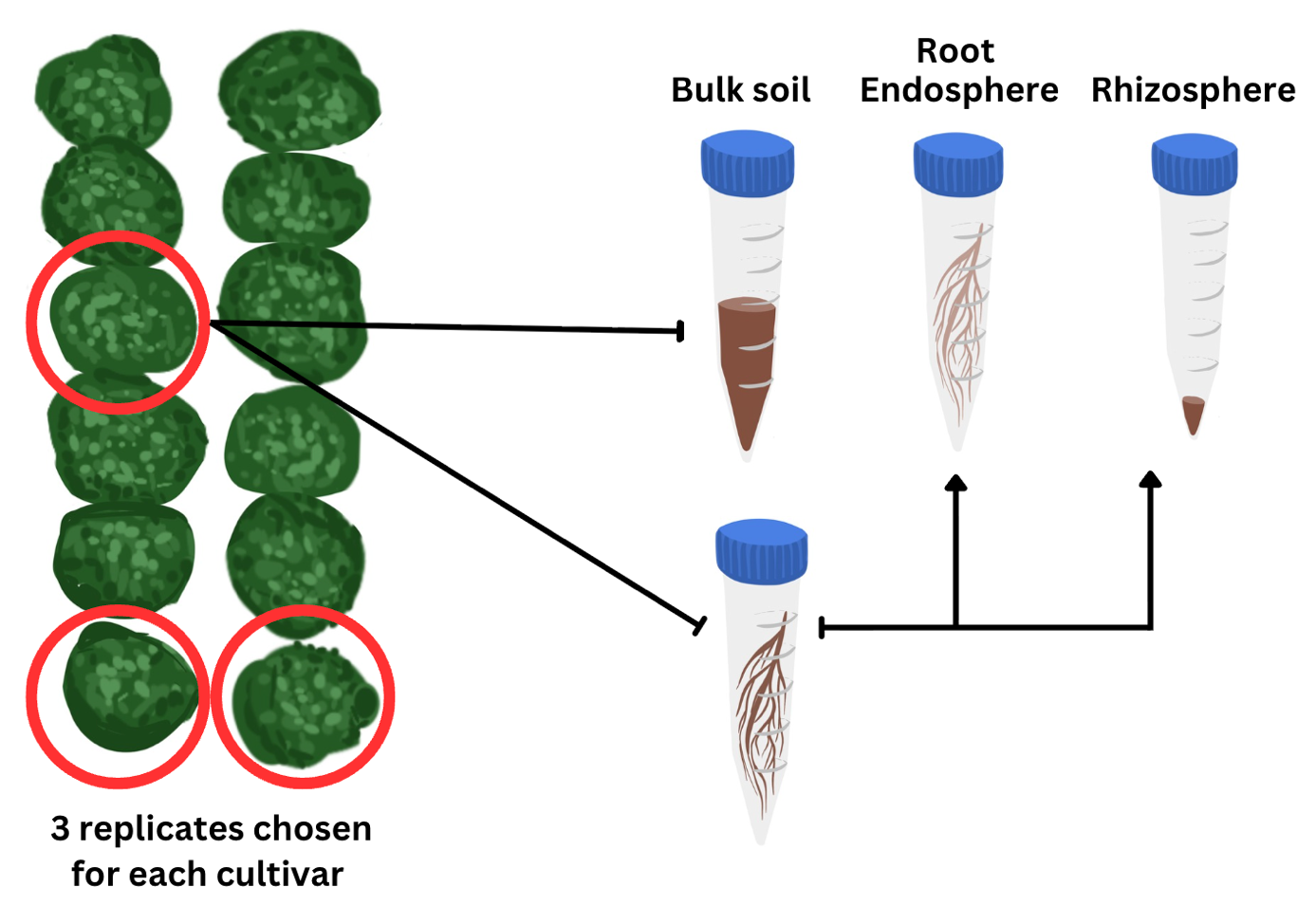

Fig S3: Drawn schematic of sampling procedure for each genotype. 3 replicates were taken from the same location in each genotype plot (exact plant locations circled in red). Samples were separated into two tubes: one containing bulk soil and anothercontaining root cutting that were then washed to separate rhizosphere soil from the root tissue.

### *Table S1.1. Bulk soil Kruskal-Wallis results for alpha diversity metrics of bacterial (16S) community*

| **Alpha Diversity Metric** | **Grouping Variable** | **Kruskal-Wallis χ²** | **p-value** |
| --- | --- | --- | --- |
| Observed OTUs | Host Phenotype | 15.252 | < 0.001 |
| Chao1 | Host Phenotype | 15.252 | < 0.001 |
| ACE | Host Phenotype | 15.189 | < 0.001 |
| Shannon | Host Phenotype | 15.158 | < 0.001 |
| Simpson | Host Phenotype | 13.930 | < 0.001 |
| Fisher's α | Host Phenotype | 15.252 | < 0.001 |
| Observed OTUs | Host Genotype | 16.229 | 0.006 |
| Chao1 | Host Genotype | 16.229 | 0.006 |
| ACE | Host Genotype | 16.144 | 0.006 |
| Shannon | Host Genotype | 15.924 | 0.007 |
| Simpson | Host Genotype | 14.731 | 0.012 |
| Fisher's α | Host Genotype | 16.229 | 0.006 |

### *Table S1.2. Dunn Post-hoc pairwise comparisons with BH correction of alpha diversity metrics for bacterial (16S) communities by Host Phenotype*

| **Pairwise Comparison** | **Z statistic** | **Unadjusted p** | **Adjusted p (BH)** | **Alpha Diversity Metric** | **Grouping** |
| --- | --- | --- | --- | --- | --- |
| Sensitive vs Water saver | -1.952712 | 0.05085373226 | 0.0508537323 | Observed OTUs | Host Phenotype |
| Sensitive vs Water spender | -3.905424 | 0.00009406033 | 0.0002821810 | Observed OTUs | Host Phenotype |
| Water saver vs Water spender | -1.952712 | 0.05085373226 | 0.0762805984 | Observed OTUs | Host Phenotype |
| Sensitive vs Water saver | -1.952712 | 0.05085373226 | 0.0508537323 | Chao1 | Host Phenotype |
| Sensitive vs Water spender | -3.905424 | 0.00009406033 | 0.0002821810 | Chao1 | Host Phenotype |
| Water saver vs Water spender | -1.952712 | 0.05085373226 | 0.0762805984 | Chao1 | Host Phenotype |
| Sensitive vs Water saver | -1.948669 | 0.05133495388 | 0.0513349539 | ACE | Host Phenotype |
| Sensitive vs Water spender | -3.897338 | 0.00009725576 | 0.0002917673 | ACE | Host Phenotype |
| Water saver vs Water spender | -1.948669 | 0.05133495388 | 0.0770024308 | ACE | Host Phenotype |
| Sensitive vs Water saver | -1.946657 | 0.05157586362 | 0.0515758636 | Shannon | Host Phenotype |
| Sensitive vs Water spender | -3.893314 | 0.00009888398 | 0.0002966519 | Shannon | Host Phenotype |
| Water saver vs Water spender | -1.946657 | 0.05157586362 | 0.0773637954 | Shannon | Host Phenotype |
| Sensitive vs Water saver | -1.784436 | 0.07435290537 | 0.0743529054 | Simpson | Host Phenotype |
| Sensitive vs Water spender | -3.731093 | 0.00019065106 | 0.0005719532 | Simpson | Host Phenotype |
| Water saver vs Water spender | -1.946657 | 0.05157586362 | 0.0773637954 | Simpson | Host Phenotype |
| Sensitive vs Water saver | -1.952712 | 0.05085373226 | 0.0508537323 | Fisher's α | Host Phenotype |
| Sensitive vs Water spender | -3.905424 | 0.00009406033 | 0.0002821810 | Fisher's α | Host Phenotype |
| Water saver vs Water spender | -1.952712 | 0.05085373226 | 0.0762805984 | Fisher's α | Host Phenotype |

### *Table S1.3. Dunn Post-hoc pairwise comparisons with BH correction of alpha diversity metrics for bacterial (16S) communities by Host Genotype. Names are abbreviated for clarity.*

| **Pairwise Comparison** | **Z statistic** | **Unadjusted p** | **Adjusted p (BH)** | **Alpha Diversity Metric** | **Grouping** |
| --- | --- | --- | --- | --- | --- |
| AP3 vs AU-16-28 | -1.11229169 | 0.2660127563 | 0.399019134 | Observed OTUs | Host Genotype |
| AP3 vs AU-17 | -2.07116383 | 0.0383434881 | 0.143788080 | Observed OTUs | Host Genotype |
| AU-16-28 vs AU-17 | -0.95887215 | 0.3376231588 | 0.422028948 | Observed OTUs | Host Genotype |
| AP3 vs Line 8 | -0.95887215 | 0.3376231588 | 0.460395216 | Observed OTUs | Host Genotype |
| AU-16-28 vs Line 8 | 0.15341954 | 0.8780674341 | 0.878067434 | Observed OTUs | Host Genotype |
| AU-17 vs Line 8 | 1.11229169 | 0.2660127563 | 0.443354594 | Observed OTUs | Host Genotype |
| AP3 vs PI-39 | 0.69038794 | 0.4899502559 | 0.524946703 | Observed OTUs | Host Genotype |
| AU-16-28 vs PI-39 | 1.80267963 | 0.0714385426 | 0.178596356 | Observed OTUs | Host Genotype |
| AU-17 vs PI-39 | 2.76155178 | 0.0057527394 | 0.028763697 | Observed OTUs | Host Genotype |
| Line 8 vs PI-39 | 1.64926009 | 0.0990943622 | 0.185801929 | Observed OTUs | Host Genotype |
| AP3 vs PI-50 | -2.76155178 | 0.0057527394 | 0.043145546 | Observed OTUs | Host Genotype |
| AU-16-28 vs PI-50 | -1.64926009 | 0.0990943622 | 0.212345062 | Observed OTUs | Host Genotype |
| AU-17 vs PI-50 | -0.69038794 | 0.4899502559 | 0.565327218 | Observed OTUs | Host Genotype |
| Line 8 vs PI-50 | -1.80267963 | 0.0714385426 | 0.214315628 | Observed OTUs | Host Genotype |
| PI-39 vs PI-50 | -3.45193972 | 0.0005565720 | 0.008348581 | Observed OTUs | Host Genotype |
| AP3 vs AU-16-28 | -1.11229169 | 0.2660127563 | 0.399019134 | Chao1 | Host Genotype |
| AP3 vs AU-17 | -2.07116383 | 0.0383434881 | 0.143788080 | Chao1 | Host Genotype |
| AU-16-28 vs AU-17 | -0.95887215 | 0.3376231588 | 0.422028948 | Chao1 | Host Genotype |
| AP3 vs Line 8 | -0.95887215 | 0.3376231588 | 0.460395216 | Chao1 | Host Genotype |
| AU-16-28 vs Line 8 | 0.15341954 | 0.8780674341 | 0.878067434 | Chao1 | Host Genotype |
| AU-17 vs Line 8 | 1.11229169 | 0.2660127563 | 0.443354594 | Chao1 | Host Genotype |
| AP3 vs PI-39 | 0.69038794 | 0.4899502559 | 0.524946703 | Chao1 | Host Genotype |
| AU-16-28 vs PI-39 | 1.80267963 | 0.0714385426 | 0.178596356 | Chao1 | Host Genotype |
| AU-17 vs PI-39 | 2.76155178 | 0.0057527394 | 0.028763697 | Chao1 | Host Genotype |
| Line 8 vs PI-39 | 1.64926009 | 0.0990943622 | 0.185801929 | Chao1 | Host Genotype |
| AP3 vs PI-50 | -2.76155178 | 0.0057527394 | 0.043145546 | Chao1 | Host Genotype |
| AU-16-28 vs PI-50 | -1.64926009 | 0.0990943622 | 0.212345062 | Chao1 | Host Genotype |
| AU-17 vs PI-50 | -0.69038794 | 0.4899502559 | 0.565327218 | Chao1 | Host Genotype |
| Line 8 vs PI-50 | -1.80267963 | 0.0714385426 | 0.214315628 | Chao1 | Host Genotype |
| PI-39 vs PI-50 | -3.45193972 | 0.0005565720 | 0.008348581 | Chao1 | Host Genotype |
| AP3 vs AU-16-28 | -0.99516238 | 0.3196572971 | 0.399571621 | ACE | Host Genotype |
| AP3 vs AU-17 | -2.06687571 | 0.0387458675 | 0.145297003 | ACE | Host Genotype |
| AU-16-28 vs AU-17 | -1.07171333 | 0.2838488123 | 0.425773218 | ACE | Host Genotype |
| AP3 vs Line 8 | -1.07171333 | 0.2838488123 | 0.473081354 | ACE | Host Genotype |
| AU-16-28 vs Line 8 | -0.07655095 | 0.9389807790 | 0.938980779 | ACE | Host Genotype |
| AU-17 vs Line 8 | 0.99516238 | 0.3196572971 | 0.435896314 | ACE | Host Genotype |
| AP3 vs PI-39 | 0.68895857 | 0.4908493407 | 0.525910008 | ACE | Host Genotype |
| AU-16-28 vs PI-39 | 1.68412095 | 0.0921582973 | 0.172796807 | ACE | Host Genotype |
| AU-17 vs PI-39 | 2.75583427 | 0.0058542651 | 0.029271325 | ACE | Host Genotype |
| Line-8 vs PI-39 | 1.76067190 | 0.0782939519 | 0.195734880 | ACE | Host Genotype |
| AP3 vs PI-50 | -2.75583427 | 0.0058542651 | 0.043906988 | ACE | Host Genotype |
| AU-16-28 vs PI-50 | -1.76067190 | 0.0782939519 | 0.234881856 | ACE | Host Genotype |
| AU-17 vs PI-50 | -0.68895857 | 0.4908493407 | 0.566364624 | ACE | Host Genotype |
| Line 8 vs PI-50 | -1.68412095 | 0.0921582973 | 0.197482066 | ACE | Host Genotype |
| PI-39 vs PI-50 | -3.44479284 | 0.0005714972 | 0.008572458 | ACE | Host Genotype |
| AP3 vs AU-16-28 | -0.99413485 | 0.3201572224 | 0.400196528 | Shannon | Host Genotype |
| AP3 vs AU-17 | -2.14121352 | 0.0322568236 | 0.120963088 | Shannon | Host Genotype |
| AU-16-28 vs AU-17 | -1.14707867 | 0.2513491088 | 0.418915181 | Shannon | Host Genotype |
| AP3 vs Line 8 | -1.07060676 | 0.2843462835 | 0.387744932 | Shannon | Host Genotype |
| AU-16-28 vs Line 8 | -0.07647191 | 0.9390436601 | 0.939043660 | Shannon | Host Genotype |
| AU-17 vs Line 8 | 1.07060676 | 0.2843462835 | 0.426519425 | Shannon | Host Genotype |
| AP3 vs PI-39 | 0.68824720 | 0.4912971242 | 0.566881297 | Shannon | Host Genotype |
| AU-16-28 vs PI-39 | 1.68238205 | 0.0924947801 | 0.198203100 | Shannon | Host Genotype |
| AU-17 vs PI-39 | 2.82946072 | 0.0046626524 | 0.034969893 | Shannon | Host Genotype |
| Line 8 vs PI-39 | 1.75885396 | 0.0786023169 | 0.235806951 | Shannon | Host Genotype |
| AP3 vs PI-50 | -2.67651690 | 0.0074391815 | 0.037195907 | Shannon | Host Genotype |
| AU-16-28 vs PI-50 | -1.68238205 | 0.0924947801 | 0.231236950 | Shannon | Host Genotype |
| AU-17 vs PI-50 | -0.53530338 | 0.5924400902 | 0.634757239 | Shannon | Host Genotype |
| Line 8 vs PI-50 | -1.60591014 | 0.1082936559 | 0.203050605 | Shannon | Host Genotype |
| PI-39 vs PI-50 | -3.36476410 | 0.0007660913 | 0.011491369 | Shannon | Host Genotype |
| AP3 vs AU-16-28 | -0.84119102 | 0.4002409281 | 0.461816455 | Simpson | Host Genotype |
| AP3 vs AU-17 | -2.06474160 | 0.0389474557 | 0.146052959 | Simpson | Host Genotype |
| AU-16-28 vs AU-17 | -1.22355058 | 0.2211218122 | 0.331682718 | Simpson | Host Genotype |
| AP3 vs Line 8 | -0.84119102 | 0.4002409281 | 0.500301160 | Simpson | Host Genotype |
| AU-16-28 vs Line 8 | 0.00000000 | 1.0000000000 | 1.000000000 | Simpson | Host Genotype |
| AU-17 vs Line 8 | 1.22355058 | 0.2211218122 | 0.368536354 | Simpson | Host Genotype |
| AP3 vs PI-39 | 0.84119102 | 0.4002409281 | 0.545783084 | Simpson | Host Genotype |
| AU-16-28 vs PI-39 | 1.68238205 | 0.0924947801 | 0.231236950 | Simpson | Host Genotype |
| AU-17 vs PI-39 | 2.90593263 | 0.0036616028 | 0.027462021 | Simpson | Host Genotype |
| Line 8 vs PI-39 | 1.68238205 | 0.0924947801 | 0.277484340 | Simpson | Host Genotype |
| AP3 vs PI-50 | -2.37062925 | 0.0177578339 | 0.088789169 | Simpson | Host Genotype |
| AU-16-28 vs PI-50 | -1.52943823 | 0.1261558422 | 0.236542204 | Simpson | Host Genotype |
| AU-17 vs PI-50 | -0.30588765 | 0.7596901931 | 0.813953778 | Simpson | Host Genotype |
| Line 8 vs PI-50 | -1.52943823 | 0.1261558422 | 0.270333948 | Simpson | Host Genotype |
| PI-39 vs PI-50 | -3.21182027 | 0.0013189686 | 0.019784529 | Simpson | Host Genotype |
| AP3 vs AU-16-28 | -1.11229169 | 0.2660127563 | 0.399019134 | Fisher's α | Host Genotype |
| AP3 vs AU-17 | -2.07116383 | 0.0383434881 | 0.143788080 | Fisher's α | Host Genotype |
| AU-16-28 vs AU-17 | -0.95887215 | 0.3376231588 | 0.422028948 | Fisher's α | Host Genotype |
| AP3 vs Line 8 | -0.95887215 | 0.3376231588 | 0.460395216 | Fisher's α | Host Genotype |
| AU-16-28 vs Line 8 | 0.15341954 | 0.8780674341 | 0.878067434 | Fisher's α | Host Genotype |
| AU-17 vs Line 8 | 1.11229169 | 0.2660127563 | 0.443354594 | Fisher's α | Host Genotype |
| AP3 vs PI-39 | 0.69038794 | 0.4899502559 | 0.524946703 | Fisher's α | Host Genotype |
| AU-16-28 vs PI-39 | 1.80267963 | 0.0714385426 | 0.178596356 | Fisher's α | Host Genotype |
| AU-17 vs PI-39 | 2.76155178 | 0.0057527394 | 0.028763697 | Fisher's α | Host Genotype |
| Line 8 vs PI-39 | 1.64926009 | 0.0990943622 | 0.185801929 | Fisher's α | Host Genotype |
| AP3 vs PI-50 | -2.76155178 | 0.0057527394 | 0.043145546 | Fisher's α | Host Genotype |
| AU-16-28 vs PI-50 | -1.64926009 | 0.0990943622 | 0.212345062 | Fisher's α | Host Genotype |
| AU-17 vs PI-50 | -0.69038794 | 0.4899502559 | 0.565327218 | Fisher's α | Host Genotype |
| Line 8 vs PI-50 | -1.80267963 | 0.0714385426 | 0.214315628 | Fisher's α | Host Genotype |
| PI-39 vs PI-50 | -3.45193972 | 0.0005565720 | 0.008348581 | Fisher's α | Host Genotype |

Table S2.1. Rhizosphere soil Kruskal-Wallis results for alpha diversity metrics of bacterial (16S) community.

| **Alpha Diversity Metric** | **Grouping Variable** | **Kruskal-Wallis χ²** | **p-value** |
| --- | --- | --- | --- |
| Observed OTUs | Host Phenotype | 5.873 | 0.053 |
| Chao1 | Host Phenotype | 5.918 | 0.052 |
| ACE | Host Phenotype | 5.400 | 0.067 |
| Shannon | Host Phenotype | 5.474 | 0.065 |
| Simpson | Host Phenotype | 5.099 | 0.078 |
| Fisher's α | Host Phenotype | 5.873 | 0.053 |
| Observed OTUs | Host Genotype | 7.917 | 0.161 |
| Chao1 | Host Genotype | 8.248 | 0.143 |
| ACE | Host Genotype | 7.606 | 0.179 |
| Shannon | Host Genotype | 9.140 | 0.104 |
| Simpson | Host Genotype | 7.690 | 0.174 |
| Fisher's α | Host Genotype | 7.917 | 0.161 |

Table 3.1: Root endosphere Kruskal-Wallis results for alpha diversity metrics of bacterial (16S) community

| **Alpha Diversity Metric** | **Grouping Variable** | **Kruskal-Wallis χ²** | **p-value** |
| --- | --- | --- | --- |
| Observed OTUs | Host Phenotype | 2.754 | 0.252 |
| Chao1 | Host Phenotype | 2.754 | 0.252 |
| ACE | Host Phenotype | 1.611 | 0.447 |
| Shannon | Host Phenotype | 1.256 | 0.534 |
| Simpson | Host Phenotype | 1.027 | 0.598 |
| Fisher's α | Host Phenotype | 2.754 | 0.252 |
| Observed OTUs | Host Genotype | 4.658 | 0.459 |
| Chao1 | Host Genotype | 4.658 | 0.459 |
| ACE | Host Genotype | 5.687 | 0.338 |
| Shannon | Host Genotype | 3.942 | 0.558 |
| Simpson | Host Genotype | 3.775 | 0.582 |
| Fisher's α | Host Genotype | 4.658 | 0.459 |

Table S4.1: Bulk soil Kruskal-Wallis results for alpha diversity metrics of fungal (ITS) community.

| **Alpha Diversity Metric** | **Grouping Variable** | **Kruskal-Wallis χ²** | **p-value** |
| --- | --- | --- | --- |
| Observed OTUs | Host Phenotype | 2.004 | 0.367 |
| Chao1 | Host Phenotype | 2.854 | 0.240 |
| ACE | Host Phenotype | 2.561 | 0.278 |
| Shannon | Host Phenotype | 0.105 | 0.949 |
| Simpson | Host Phenotype | 1.626 | 0.444 |
| Fisher's α | Host Phenotype | 2.004 | 0.367 |
| Observed OTUs | Host Genotype | 7.800 | 0.168 |
| Chao1 | Host Genotype | 7.222 | 0.205 |
| ACE | Host Genotype | 7.737 | 0.171 |
| Shannon | Host Genotype | 10.497 | 0.062 |
| Simpson | Host Genotype | 11.480 | 0.043 |
| Fisher's α | Host Genotype | 7.800 | 0.168 |

Table S4.2: Dunn Post-hoc pairwise comparisons with BH correction of alpha diversity metrics for fungal (ITS) bulk soil communities by Host Genotype. Names are abbreviated for clarity.

| **Pairwise Comparison** | **Z statistic** | **Unadjusted p** | **Adjusted p (BH)** | **Alpha Diversity Metric** | **Grouping** |
| --- | --- | --- | --- | --- | --- |
| AP3 vs AU-16-28 | -0.9176629 | 0.358795358 | 0.53819304 | Simpson | Host Genotype |
| AP3 vs AU-17 | -0.3823596 | 0.702194685 | 0.75235145 | Simpson | Host Genotype |
| AU-16-28 vs AU-17 | 0.5353034 | 0.592440090 | 0.68358472 | Simpson | Host Genotype |
| AP3 vs Line 8 | -2.7529888 | 0.005905392 | 0.08858087 | Simpson | Host Genotype |
| AU-16-28 vs Line 8 | -1.8353259 | 0.066457420 | 0.19937226 | Simpson | Host Genotype |
| AU-17 vs Line 8 | -2.3706292 | 0.017757834 | 0.13318375 | Simpson | Host Genotype |
| AP3 vs PI-39 | -1.9117978 | 0.055902136 | 0.20963301 | Simpson | Host Genotype |
| AU-16-28 vs PI-39 | -0.9941348 | 0.320157222 | 0.53359537 | Simpson | Host Genotype |
| AU-17 vs PI-39 | -1.5294382 | 0.126155842 | 0.27033395 | Simpson | Host Genotype |
| Line 8 vs PI-39 | 0.8411910 | 0.400240928 | 0.54578308 | Simpson | Host Genotype |
| AP3 vs PI-50 | -2.0647416 | 0.038947456 | 0.19473728 | Simpson | Host Genotype |
| AU-16-28 vs PI-50 | -1.1470787 | 0.251349109 | 0.47127958 | Simpson | Host Genotype |
| AU-17 vs PI-50 | -1.6823820 | 0.092494780 | 0.23123695 | Simpson | Host Genotype |
| Line 8 vs PI-50 | 0.6882472 | 0.491297124 | 0.61412141 | Simpson | Host Genotype |
| PI-39 vs PI-50 | -0.1529438 | 0.878442577 | 0.87844258 | Simpson | Host Genotype |

Table S5.1: Rhizosphere soil Kruskal-Wallis results for alpha diversity metrics of fungal (ITS) community.

| **Alpha Diversity Metric** | **Grouping Variable** | **Kruskal-Wallis χ²** | **p-value** |
| --- | --- | --- | --- |
| Observed OTUs | Host Phenotype | 1.560 | 0.458 |
| Chao1 | Host Phenotype | 0.737 | 0.692 |
| ACE | Host Phenotype | 1.485 | 0.476 |
| **Shannon** | **Host Phenotype** | **7.614** | **0.022** |
| Simpson | Host Phenotype | 8.561 | 0.014 |
| Fisher's α | Host Phenotype | 1.560 | 0.458 |
| Observed OTUs | Host Genotype | 4.904 | 0.428 |
| Chao1 | Host Genotype | 3.620 | 0.605 |
| ACE | Host Genotype | 3.971 | 0.554 |
| **Shannon** | **Host Genotype** | **11.246** | **0.047** |
| Simpson | Host Genotype | 11.339 | 0.045 |
| Fisher's α | Host Genotype | 4.904 | 0.428 |

Table S5.2: Dunn Post-hoc pairwise comparisons with BH correction of alpha diversity metrics for fungal (ITS) rhizosphere soil communities by host phenotype.

| **Pairwise Comparison** | **Z statistic** | **Unadjusted p** | **Adjusted p (BH)** | **Alpha Diversity Metric** | **Grouping** |
| --- | --- | --- | --- | --- | --- |
| Sensitive vs Water saver | 1.297771 | 0.194365911 | 0.19436591 | Shannon | Host Phenotype |
| Sensitive vs Water spender | -1.459993 | 0.144292055 | 0.21643808 | Shannon | Host Phenotype |
| Water saver vs Water spender | -2.757764 | 0.005819817 | 0.01745945 | Shannon | Host Phenotype |
| Sensitive vs Water saver | 1.622214 | 0.104757490 | 0.15713623 | Simpson | Host Phenotype |
| Sensitive vs Water spender | -1.297771 | 0.194365911 | 0.19436591 | Simpson | Host Phenotype |
| Water saver vs Water spender | -2.919986 | 0.003500476 | 0.01050143 | Simpson | Host Phenotype |

Table S5.3: Dunn Post-hoc pairwise comparisons with BH correction of alpha diversity metrics for fungal (ITS) rhizosphere soil communities by host genotype. Names are abbreviated for clarity.

| **Pairwise Comparison** | **Z statistic** | **Unadjusted p** | **Adjusted p (BH)** | **Alpha Diversity Metric** | **Grouping** |
| --- | --- | --- | --- | --- | --- |
| AP3 vs AU-16-28 | 0.99413485 | 0.320157222 | 0.48023583 | Shannon | Host Genotype |
| AP3 vs AU-17 | 0.07647191 | 0.939043660 | 0.93904366 | Shannon | Host Genotype |
| AU-16-28 vs AU-17 | -0.91766294 | 0.358795358 | 0.44849420 | Shannon | Host Genotype |
| AP3 vs Line 8 | 2.06474160 | 0.038947456 | 0.19473728 | Shannon | Host Genotype |
| AU-16-28 vs Line 8 | 1.07060676 | 0.284346284 | 0.53314928 | Shannon | Host Genotype |
| AU-17 vs Line 8 | 1.98826969 | 0.046781871 | 0.17543202 | Shannon | Host Genotype |
| AP3 vs PI-39 | 1.22355058 | 0.221121812 | 0.55280453 | Shannon | Host Genotype |
| AU-16-28 vs PI-39 | 0.22941573 | 0.818545808 | 0.87701337 | Shannon | Host Genotype |
| AU-17 vs PI-39 | 1.14707867 | 0.251349109 | 0.53860523 | Shannon | Host Genotype |
| Line 8 vs PI-39 | -0.84119102 | 0.400240928 | 0.46181646 | Shannon | Host Genotype |
| AP3 vs PI-50 | -0.91766294 | 0.358795358 | 0.48926640 | Shannon | Host Genotype |
| AU-16-28 vs PI-50 | -1.91179778 | 0.055902136 | 0.16770641 | Shannon | Host Genotype |
| AU-17 vs PI-50 | -0.99413485 | 0.320157222 | 0.53359537 | Shannon | Host Genotype |
| Line 8 vs PI-50 | -2.98240454 | 0.002859938 | 0.04289907 | Shannon | Host Genotype |
| PI-39 vs PI-50 | -2.14121352 | 0.032256824 | 0.24192618 | Shannon | Host Genotype |
| AP3 vs AU-16-28 | 1.22355058 | 0.221121812 | 0.47383245 | Simpson | Host Genotype |
| AP3 vs AU-17 | 0.00000000 | 1.000000000 | 1.00000000 | Simpson | Host Genotype |
| AU-16-28 vs AU-17 | -1.22355058 | 0.221121812 | 0.55280453 | Simpson | Host Genotype |
| AP3 vs Line 8 | 2.21768543 | 0.026576289 | 0.13288144 | Simpson | Host Genotype |
| AU-16-28 vs Line 8 | 0.99413485 | 0.320157222 | 0.43657803 | Simpson | Host Genotype |
| AU-17 vs Line 8 | 2.21768543 | 0.026576289 | 0.19932216 | Simpson | Host Genotype |
| AP3 vs PI-39 | 1.14707867 | 0.251349109 | 0.41891518 | Simpson | Host Genotype |
| AU-16-28 vs PI-39 | -0.07647191 | 0.939043660 | 1.00000000 | Simpson | Host Genotype |
| AU-17 vs PI-39 | 1.14707867 | 0.251349109 | 0.47127958 | Simpson | Host Genotype |
| Line 8 vs PI-39 | -1.07060676 | 0.284346284 | 0.42651943 | Simpson | Host Genotype |
| AP3 vs PI-50 | -0.68824720 | 0.491297124 | 0.56688130 | Simpson | Host Genotype |
| AU-16-28 vs PI-50 | -1.91179778 | 0.055902136 | 0.20963301 | Simpson | Host Genotype |
| AU-17 vs PI-50 | -0.68824720 | 0.491297124 | 0.61412141 | Simpson | Host Genotype |
| Line 8 vs PI-50 | -2.90593263 | 0.003661603 | 0.05492404 | Simpson | Host Genotype |
| PI-39 vs PI-50 | -1.83532587 | 0.066457420 | 0.19937226 | Simpson | Host Genotype |

Table S6.1: Root endosphere Kruskal-Wallis results for alpha diversity metrics of fungal (ITS) community.

| **Alpha Diversity Metric** | **Grouping Variable** | **Kruskal-Wallis χ²** | **p-value** |
| --- | --- | --- | --- |
| Observed OTUs | Host Phenotype | 3.690 | 0.158 |
| Chao1 | Host Phenotype | 3.134 | 0.209 |
| ACE | Host Phenotype | 5.206 | 0.074 |
| **Shannon** | **Host Phenotype** | **6.899** | **0.018** |
| Simpson | Host Phenotype | 5.206 | 0.074 |
| Fisher's α | Host Phenotype | 3.690 | 0.158 |
| Observed OTUs | Host Genotype | 8.162 | 0.148 |
| Chao1 | Host Genotype | 7.832 | 0.166 |
| ACE | Host Genotype | 8.441 | 0.134 |
| Shannon | Host Genotype | 9.382 | 0.095 |
| Simpson | Host Genotype | 9.618 | 0.087 |
| Fisher's α | Host Genotype | 8.162 | 0.148 |

Table S6.2: Dunn Post-hoc pairwise comparisons with BH correction of alpha diversity metrics for fungal (ITS) root endosphere communities by host phenotype.

| **Pairwise Comparison** | **Z statistic** | **Unadjusted p** | **Adjusted p (BH)** | **Alpha Diversity Metric** | **Grouping** |
| --- | --- | --- | --- | --- | --- |
| Sensitive vs Water saver | -0.9828067 | 0.325702563 | 0.32570256 | Shannon | Host Phenotype |
| Sensitive vs Water spender | -2.5904235 | 0.009585791 | 0.02875737 | Shannon | Host Phenotype |
| Water saver vs Water spender | -1.7228024 | 0.084924264 | 0.12738640 | Shannon | Host Phenotype |

### *Table S7.1. Bacterial (16S) Bulk Soil Beta Diversity Results for Bray-Curtis and Unweighted Unifrac distance metrics.*

| **Distance** | **Grouping** | | **Pseudo-F** | | **R²** | **PERMANOVA p** | **Betadisper p** |
| --- | --- | --- | --- | --- | --- | --- | --- |
| Bray | Host Phenotype | 1.690031 | | 0.1838983 | | **0.009** | 0.837 |
| Bray | Host Genotype | 1.441981 | | 0.3753223 | | **0.011** | 0.849 |
| Unifrac | Host Phenotype | 2.456868 | | 0.2467511 | | **0.005** | 0.750 |
| Unifrac | Host Genotype | 1.534284 | | 0.3899779 | | **0.045** | 0.755 |

### *Table S7.2. Bacterial (16S) Rhizosphere Soil Beta Diversity Results for Bray-Curtis and Unweighted Unifrac distance metrics.*

| **Distance** | **Grouping** | **Pseudo-F** | **R²** | **PERMANOVA p** | **Betadisper p** |
| --- | --- | --- | --- | --- | --- |
| Bray | Host Phenotype | 1.238743 | 0.1417530 | 0.116 | 0.249 |
| Bray | Host Genotype | 1.125533 | 0.3192520 | 0.179 | 0.681 |
| Unifrac | Host Phenotype | 1.096336 | 0.1275353 | 0.332 | 0.436 |
| Unifrac | Host Genotype | 1.033728 | 0.3010512 | 0.406 | 0.834 |

### *Table S7.3. Bacterial (16S) Root Endosphere Beta Diversity Results for Bray-Curtis and Unweighted Unifrac distance metrics.*

| **Distance** | | **Grouping** | **Pseudo-F** | **R²** | **PERMANOVA p** | **Betadisper p** |
| --- | --- | --- | --- | --- | --- | --- |
| Bray | Host Phenotype | | 1.0303632 | 0.1465590 | 0.403 | 0.495 |
| Bray | Host Genotype | | 0.8556061 | 0.3221886 | 0.660 | 0.559 |
| Unifrac | Host Phenotype | | 1.3057445 | 0.1787285 | 0.234 | 0.685 |
| Unifrac | Host Genotype | | 0.9274759 | 0.3400492 | 0.574 | 0.136 |

### *Table S8.1. Fungal (ITS) Bulk Soil Beta Diversity Results for Bray-Curtis and Unweighted Unifrac distance metrics.*

| **Distance** | **Grouping** | **Pseudo_F** | **R2** | **p_value** | **betadisper_p** |
| --- | --- | --- | --- | --- | --- |
| Bray | Host Phenotype | 1.931142 | 0.2047623 | 0.002 | 0.445 |
| Bray | Host Genotype | 1.750313 | 0.4217304 | <0.001 | 0.550 |
| Unifrac | Host Phenotype | 1.999313 | 0.2104692 | 0.023 | 0.157 |
| Unifrac | Host Genotype | 2.002594 | 0.4548668 | 0.006 | 0.404 |

### *Table S8.2. Fungal (ITS) Rhizosphere Soil Beta Diversity Results for Bray-Curtis and Unweighted Unifrac distance metrics.*

| **Distance** | **Grouping** | **Pseudo-F** | **R²** | **PERMANOVA p** | **Betadisper p** |
| --- | --- | --- | --- | --- | --- |
| Bray | Host Phenotype | 2.0840316 | 0.2174483 | 0.053 | 0.432 |
| Bray | Host Genotype | 1.9821391 | 0.4523223 | **0.028** | 0.828 |
| Unifrac | Host Phenotype | 0.9770789 | 0.1152613 | 0.480 | 0.960 |
| Unifrac | Host Genotype | 1.3382756 | 0.3579928 | 0.068 | 0.885 |

### *Table S8.3. Fungal (ITS) Root Endosphere Beta Diversity Results for Bray-Curtis and Unweighted Unifrac distance metrics.*

| **Distance** | **Grouping** | **Pseudo-F** | **R²** | **PERMANOVA p** | **Betadisper p** |
| --- | --- | --- | --- | --- | --- |
| Bray | Host Phenotype | 2.472137 | 0.2755349 | **0.037** | 0.877 |
| Bray | Host Genotype | 1.650538 | 0.4521356 | 0.109 | 0.795 |
| Unifrac | Host Phenotype | 1.136296 | 0.1488019 | 0.326 | 0.818 |
| Unifrac | Host Genotype | 1.554198 | 0.4372852 | 0.059 | 0.374 |

*Table S9.1. Nested PERMANOVA model results for bacterial (16S) bulk soil community.*
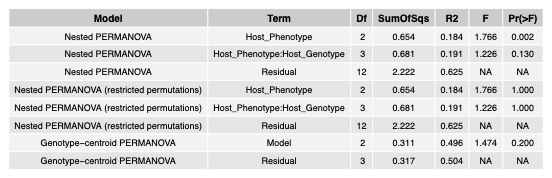

Table S9.2. *Nested PERMANOVA model results for bacterial (16S) rhizosphere soil community.*

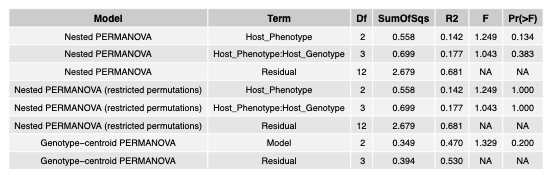

Table S9.3. *Nested PERMANOVA model results for bacterial (16S) root endosphere community.*
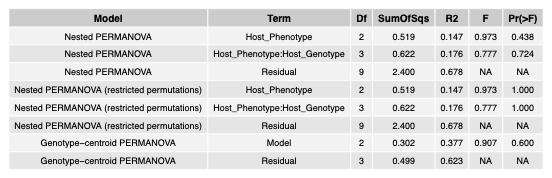

Table S9.4. *Nested PERMANOVA model results for fungal (ITS) bulk soil community.*

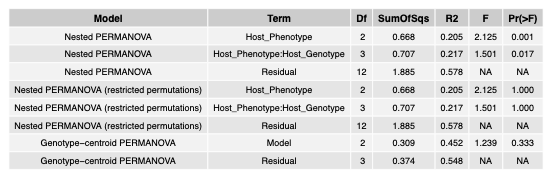

Table S9.5. *Nested PERMANOVA model results for fungal (ITS) rhizosphere soil community.*
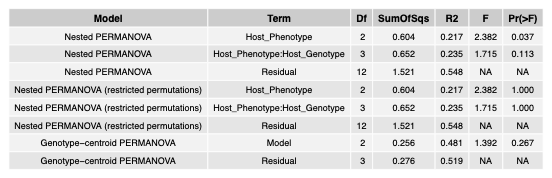

Table S9.6. *Nested PERMANOVA model results for fungal (ITS) root endosphere community.*

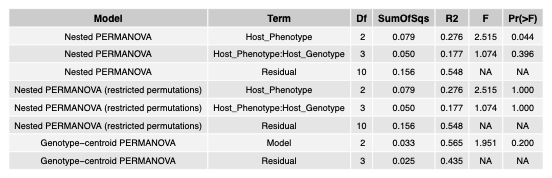

*Table S10. Sloan neutral community model fit statistics of bacterial (16S) bulk soil, rhizosphere, and root endosphere communities by host Phenotype. m, migration rate; Nm, effective number of migrants; R2, proportion of variance in occurrence frequency explained by the neutral model.*

**Bulk Soil**

| **Host Phenotype** | **n taxa** | **m** | **95% CI (m)** | **Nm** | **R²** | **Above, n (%)** | **Below, n (%)** | **Neutral, n (%)** |
| --- | --- | --- | --- | --- | --- | --- | --- | --- |
| Water spender | 93 | 0.0266 | 0.0246–0.0288 | 14.5 | 0.929 | 0 (0%) | 0 (0%) | 93 (100%) |
| Sensitive | 31 | 0.0310 | 0.026–0.0368 | 5.7 | 0.875 | 0 (0%) | 0 (0%) | 31 (100%) |
| Water saver | 59 | 0.0323 | 0.0291–0.0358 | 10.0 | 0.925 | 0 (0%) | 0 (0%) | 59 (100%) |

**Rhizosphere**

| **Host Phenotype** | **n taxa** | **m** | **95% CI (m)** | **Nm** | **R²** | **Above, n (%)** | **Below, n (%)** | **Neutral, n (%)** |
| --- | --- | --- | --- | --- | --- | --- | --- | --- |
| Water spender | 97 | 0.0190 | 0.0166–0.0217 | 11.3 | 0.753 | 0 (0%) | 1 (1%) | 96 (99%) |
| Sensitive | 71 | 0.0163 | 0.0138–0.0191 | 7.3 | 0.779 | 0 (0%) | 0 (0%) | 71 (100%) |
| Water saver | 73 | 0.0251 | 0.0228–0.0275 | 13.1 | 0.936 | 0 (0%) | 0 (0%) | 73 (100%) |

| **Host Phenotype** | **Fit type** | **n Taxa** | **Taxa** |
| --- | --- | --- | --- |
| Water spender | Suppressed (below prediction) | 1 | *Chitinophaga sp.* |

**Endosphere**

| **Host Phenotype** | **n taxa** | **m** | **95% CI (m)** | **Nm** | **R²** | **Above, n (%)** | | **Below, n (%)** | **Neutral, n (%)** |
| --- | --- | --- | --- | --- | --- | --- | --- | --- | --- |
| Water spender | 52 | 0.01890 | 0.0158–0.0224 | 15.2 | 0.804 | 0 (0%) | | 0 (0%) | 52 (100%) |
| Sensitive | 33 | 0.00699 | 0.00442–0.0105 | 6.5 | -0.054 | 0 (0%) | | 1 (3%) | 32 (97%) |
| Water saver | 45 | 0.00696 | 0.00455–0.0101 | 5.6 | 0.277 | 0 (0%) | 2 (4.4%) | | 43 (95.6%) |

| **Host Phenotype** | **Fit type** | **n Taxa** | **Taxa** |
| --- | --- | --- | --- |
| Sensitive | Suppressed (below prediction) | 1 | *Amycolatopsis sp.* |
| Water saver | Suppressed (below prediction) | 2 | *Amycolatopsis sp., unclassified Enterobacteriaceae* |

*Table S11. Sloan neutral community model fit statistics of fungal (ITS) bulk soil, rhizosphere, and root endosphere communities by host Phenotype. m, migration rate; Nm, effective number of migrants; R2, proportion of variance in occurrence frequency explained by the neutral model.*

**Bulk Soil**

| **Host Phenotype** | **n taxa** | **m** | **95% CI (m)** | **Nm** | **R²** | **Above, n (%)** | **Below, n (%)** | **Neutral, n (%)** |
| --- | --- | --- | --- | --- | --- | --- | --- | --- |
| Water spender | 172 | 0.0562 | 0.0434–0.0729 | 96.4 | 0.370 | 3 (1.7%) | 10 (5.8%) | 159 (92.4%) |
| Sensitive | 177 | 0.1640 | 0.124–0.222 | 298.2 | 0.495 | 4 (2.3%) | 15 (8.5%) | 158 (89.3%) |
| Water saver | 165 | 0.0819 | 0.0638–0.105 | 139.0 | 0.376 | 1 (0.6%) | 14 (8.5%) | 150 (90.9%) |

| **Host Phenotype** | | **Fit type** | **n Taxa** | | **Taxa** |
| --- | --- | --- | --- | --- | --- |
| Sensitive | Enriched (above prediction) | | | 4 | *g__Rhizopus s__arrhizus, g__Sporobolomyces s__patagonicus, g__Torula sp., unclassified k__Fungi* |
| Sensitive | Suppressed (below prediction) | | | 15 | *g__Allophoma s__tropica, g__Alternaria sp., g__Bipolaris s__drechsleri, g__Colletotrichum sp., g__Minimedusa sp., g__Mortierella sp., g__Poaceascoma sp., g__Rhizophlyctis sp., g__Sistotrema s__adnatum, unclassified c__Agaricomycetes, unclassified k__Fungi, unclassified p__Ascomycota* |
| Water saver | Enriched (above prediction) | | | 1 | *unclassified k__Fungi* |
| Water saver | Suppressed (below prediction) | | | 14 | *g__Alternaria sp., g__Aspergillus s__flavus, g__Aspergillus s__ibericus, g__Cladosporium s__herbarum, g__Colletotrichum sp., g__Leucocoprinus sp., g__Linnemannia sp., g__Pseudopithomyces s__rosae, g__Sistotrema s__adnatum, unclassified f__Agaricaceae, unclassified p__Ascomycota* |
| Water spender | Enriched (above prediction) | | | 3 | *g__Rhizopus s__arrhizus, g__Spizellomycetales_gen_Incertae_sedis sp., g__Torula sp.* |
| Water spender | Suppressed (below prediction) | | | 10 | *g__Agrocybe s__retigera, g__Aspergillus s__flavus, g__Colletotrichum sp., g__Fusarium s__brevicaudatum, g__Leucocoprinus sp., g__Mortierella s__rishikesha, g__Xepicula s__leucotricha, unclassified f__Agaricaceae, unclassified k__Fungi* |

**Rhizosphere**

| **Host Phenotype** | **n taxa** | **m** | **95% CI (m)** | **Nm** | **R²** | **Above, n (%)** | **Below, n (%)** | **Neutral, n (%)** |
| --- | --- | --- | --- | --- | --- | --- | --- | --- |
| Water spender | 133 | 0.0197 | 0.0126–0.031 | 35.9 | -0.003 | 17 (12.8%) | 6 (4.5%) | 110 (82.7%) |
| Sensitive | 106 | 0.1130 | 0.0692–0.185 | 223.3 | 0.387 | 1 (0.9%) | 10 (9.4%) | 95 (89.6%) |
| Water saver | 102 | 0.1930 | 0.121–0.311 | 386.7 | 0.353 | 1 (1%) | 11 (10.8%) | 90 (88.2%) |

| **Host Phenotype** | **Fit type** | **n Taxa** | | **Taxa** |
| --- | --- | --- | --- | --- |
| Sensitive | Enriched (above prediction) | | 1 | *unclassified p__Ascomycota* |
| Sensitive | Suppressed (below prediction) | | 10 | *g__Epicoccum s__pimprinum, g__Knufia sp., g__Linnemannia sp., g__Mortierella sp., g__Talaromyces s__subaurantiacus, unclassified f__Didymellaceae* |
| Water saver | Enriched (above prediction) | | 1 | *g__Hannaella s__oryzae* |
| Water saver | Suppressed (below prediction) | | 11 | *g__Aspergillus s__flavus, g__Epicoccum sp., g__Knufia sp., g__Leucocoprinus sp., g__Linnemannia sp., g__Marasmius s__nigrobrunneus, g__Mortierella s__rishikesha, g__Talaromyces s__subaurantiacus, g__Talaromyces sp., unclassified p__Ascomycota* |
| Water spender | Enriched (above prediction) | | 17 | *g__Allophoma s__tropica, g__Candolleomyces s__luteopallidus, g__Colletotrichum s__chlorophyti, g__Colletotrichum sp., g__Diaporthe sp., g__Hannaella s__oryzae, g__Lectera s__nordwiniana, g__Mortierella sp., g__Pseudopithomyces s__rosae, g__Ramicandelaber s__taiwanensis, g__Rhizophydiales_gen_Incertae_sedis sp., g__Sporobolomyces s__patagonicus, g__Westerdykella s__ornata, unclassified f__Glomeraceae, unclassified p__Ascomycota* |
| Water spender | Suppressed (below prediction) | | 6 | *g__Agrocybe s__retigera, g__Branch06_gen_Incertae_sedis sp., g__Marasmius s__nigrobrunneus, g__Mortierella sp., g__Talaromyces sp.* |

**Endosphere**

| **Host Phenotype** | **n taxa** | **m** | **95% CI (m)** | **Nm** | **R²** | **Above, n (%)** | **Below, n (%)** | **Neutral, n (%)** |
| --- | --- | --- | --- | --- | --- | --- | --- | --- |
| Water spender | 49 | 0.188 | 0.0845–0.403 | 376.0 | 0.009 | 1 (2%) | 4 (8.2%) | 44 (89.8%) |
| Sensitive | 30 | 0.243 | NA–NA | 503.5 | -0.398 | 0 (0%) | 4 (13.3%) | 26 (86.7%) |
| Water saver | 34 | 0.269 | 0.149–0.5 | 552.6 | 0.463 | 0 (0%) | 2 (5.9%) | 32 (94.1%) |

| **Host Phenotype** | **Fit class** | **n Taxa** | | **Taxa** |
| --- | --- | --- | --- | --- |
| Sensitive | Suppressed (below prediction) | | 4 | *g__Knufia sp., unclassified p__Ascomycota* |
| Water saver | Suppressed (below prediction) | | 2 | *unclassified p__Ascomycota* |
| Water spender | Enriched (above prediction) | | 1 | *unclassified p__Ascomycota* |
| Water spender | Suppressed (below prediction) | | 4 | *g__Epicoccum s__pimprinum, g__Lectera s__nordwiniana, unclassified f__Didymellaceae, unclassified p__Ascomycota* |
